# Gene panel selection for targeted spatial transcriptomics

**DOI:** 10.1101/2023.02.03.527053

**Authors:** Yida Zhang, Viktor Petukhov, Evan Biederstedt, Richard Que, Kun Zhang, Peter V. Kharchenko

## Abstract

Targeted spatial transcriptomics hold particular promise in analysis of complex tissues. Most such methods, however, measure only a limited panel of transcripts, which need to be selected in advance to inform on the cell types or processes being studied. A limitation of existing gene selection methods is that they rely on scRNA-seq data, ignoring platform effects between technologies. Here we describe gpsFISH, a computational method to perform gene selection through optimizing detection of known cell types. By modeling and adjusting for platform effects, gpsFISH outperforms other methods. Furthermore, gpsFISH can incorporate cell type hierarchies and custom gene preferences to accommodate diverse design requirements.

## Background

The building block of complex tissues is the diverse range of cell types [1–4]. Knowing the identity and spatial location of cells from different cell types is the key for understanding how they communicate with each other to carry out specific functions and how diseases emerge when this complex network of interactions goes awry [5–11]. Single-cell RNA sequencing (scRNA-seq) provides a powerful tool to study the identity of cell types and cell states [12–17]. However, the spatial information is lost due to cell disassociation during library preparation. Recent advances in spatial transcriptomics technologies have overcome this limitation by providing ways to quantify gene expression while keeping the spatial information of cells, leading to more comprehensive and detailed understanding of diseases and normal functions [18–23].

Based on the number of transcripts that can be probed, spatial transcriptomics technologies can be broadly categorized as (1) targeted, measuring a limited panel of transcripts; and (2) untargeted, capturing all transcripts from the transcriptome. Targeted spatial transcriptomics include in situ hybridization (ISH)-based [24–28] and most in situ sequencing (ISS)-based methods [29–32]. Untargeted spatial transcriptomics include next-generation sequencing (NGS)-based methods [33–39].

Compared to untargeted spatial transcriptomics, targeted spatial transcriptomics can achieve high sensitivity and subcellular resolution. However, their targeted nature requires a panel of genes (from a few hundred to thousand) to be selected in advance to recognize cell types or processes relevant to the tissue being studied.

Gene selection methods are used to design gene panels. They can be classified into two major categories based on their gene selection objectives. One category with an imputation-based objective aims to select genes based on their ability to capture as much of transcriptional variation in the scRNA-seq data as possible. Examples range from simply selecting highly-variable genes to more advanced methods like L1000 [40], geneBasis [41], and SCMER [42]. Specifically, L1000 identified the optimal set of ‘landmark’ transcripts that construct a reduced representations of the transcriptome. geneBasis finds genes that can yield a *k*-nearest neighbor graph that is similar to the “true” graph constructed using the entire transcriptome. SCMER aims to select genes that preserve the manifold of scRNA-seq data. Another category of gene selection method with a classification-based objective selects genes given their ability to reconstruct cell classifications or relationships. Examples range from selecting differentially expressed genes (DEGs) to more advanced methods like scGeneFit [43], RankCorr [44], and NS-Forest [45]. scGeneFit selects marker genes that jointly optimize cell type recovery using a label-aware compressive classification method. RankCorr is a rank-based one-vs-all feature selection method that selects marker genes for each cell type based on a sparsity parameter that controls the number of marker genes selected per cell type. NS-Forest is a machine learning-based marker gene selection algorithm that uses the nonlinear attributes of random forest feature selection and a binary expression scoring approach to select the minimal combination of marker genes that captures the cell type identity in scRNA-seq data. All these methods can be used to design gene panels for targeted spatial transcriptomics technologies.

A key limitation of current gene selection methods is that they select genes purely based on scRNA-seq data without considering potential differences between scRNA-seq and the target spatial transcriptomics technologies. Such platform effects include systematic differences in capture efficiency of genes between platforms caused by technology-dependent factors, including detection technique and library preparation chemistry. Platform effects have been previously noted when comparing gene expression measurements from single-cell and single-nucleus RNA-seq on the same biological sample [46]. Platform effects also exist between scRNA-seq and spatial transcriptomics technologies [47–49], posing a challenge when transferring cell type information from scRNA-seq to spatial transcriptomics technologies. When selecting gene panels using scRNA-seq data, such platform-specific distortions can lead to reduced performance of selected gene panels in the resulting spatial measurements.

Besides platform effects, there are other complications involved in gene panel selection. First, current classification-based gene selection methods [43–45] treat cell types as equally distinct. However, cell types are organized in a hierarchical manner with cell subpopulations belonging to the same broad cell type more similar to each other than subpopulations belonging to different broad cell types [50–56]. Depending on the biological questions and capabilities of the assays, a gene selection method could optimize for fidelity at lower cell type resolution, or place increased emphasis on certain subgroups of cell types. More generally, this is not only useful for selecting genes that inform on cell types but can also be extended to selecting genes for other biological entities with a hierarchical structure, e.g., gene ontology and pathways [57,58]. Second, both imputation-based and classification-based gene selection methods select genes solely based on a pre-defined objective function. However, in practice of gene panel design for targeted spatial transcriptomics, there can be other criteria contributing to the gene selection. Examples range from technical factors, such as ability to design probes for targeting certain transcripts, to biological factors such as preferences for certain pathways or marker genes commonly used in the literature. A framework that takes such orthogonal preferences into consideration can be helpful in practice.

To address these challenges, we developed gpsFISH, a classification-based gene selection method that models and adjusts for the platform effects between scRNA-seq and targeted spatial transcriptomics technologies, yielding more informative gene panels and better cell type classification compared to previously published classification-based gene selection methods. In addition, gpsFISH provides options to account for cell type hierarchy and gene-specific custom preferences during gene panel design, offering flexible and finer control of cell type granularity and gene selection for different biological questions.

## Results

### Platform effects between scRNA-seq and targeted spatial transcriptomics

Even molecule counting assays carry inherent detection biases, posing challenges for joint analysis of multiple assays, such as scRNA-seq and spatially-resolved counts [47–49]. Indeed, we observed a systematic difference of transcript detection rate across platforms (**Fig. 1A**–**D**), which distorts the resulting transcriptional profile estimates. Consequently, a panel of genes selected based on scRNA-seq that works well on distinguishing cell types may not achieve similar level of performance when measured by targeted spatial transcriptomics.

**Figure 1.**
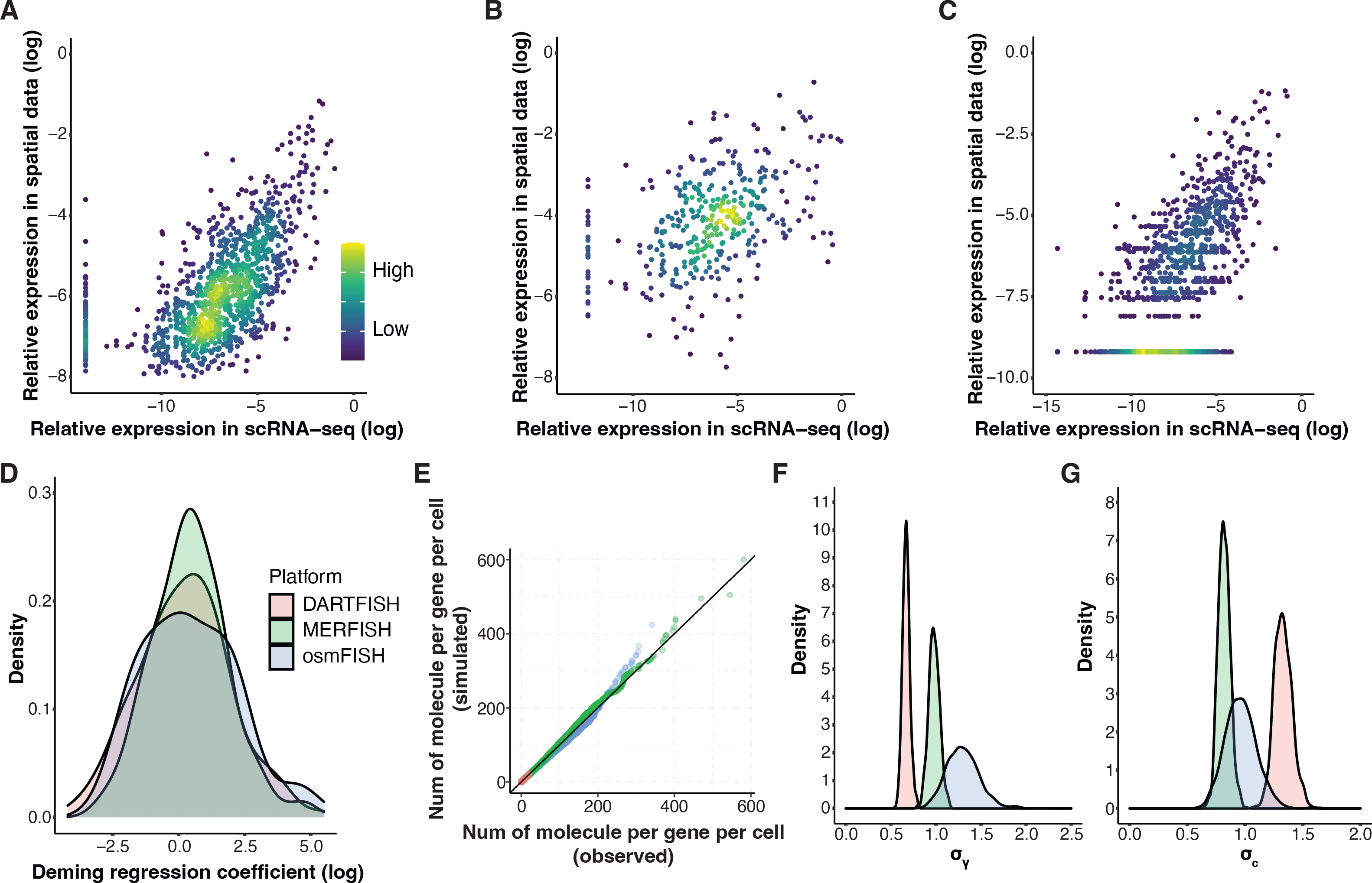
Platform effect between scRNA-seq and targeted spatial transcriptomics technologies. **A-C**: Scatter plot showing the log transformed relative expression of genes measured by both scRNA-seq and targeted spatial transcriptomics across three datasets, Moffit (**A**), Codeluppi (**B**), and Zhang (**C**), respectively. A small value is added to avoid negative infinity after log transformation. Each dot represents the relative expression of one gene in one cell type. Denominator for relative expression calculation is from all genes measured by both technologies. Color indicates density of dots. Dots should fall on the diagonal when there is no platform effect. **D**: Density plot of Deming regression coefficient for each dataset. Deming regression is fitted for each gene using relative expression measured by scRNA-seq and spatial transcriptomics data with intercept fixed to 0. **E**: Posterior predictive check of the Bayesian models fitted using each of the three datasets. QQ plot showing the distribution of simulated vs. observed spatial transcriptomics measurements. **F-G**: Density plot showing the estimated posterior distribution of σ*_y_* (**F**) and σ*_c_* (**G**).

To address this challenge, we estimate the level of gene expression distortion in targeted spatial transcriptomics data relative to scRNA-seq and from the same tissue using a Bayesian model (**Fig. S1**, **Methods**). Bayesian inference estimates the posterior distribution of distortion magnitudes, which will be used to predict the potential distortion levels for genes that have not yet been observed in a given assay. Specifically, we assume platform effects are on a per gene basis. γ*_i_* and *C_j_* represent gene specific multiplicative and additive platform effect for each gene *i*, respectively. These distortion parameters are assumed to follow two normal distributions with μ*_y_*, μ*_c_* as mean and σ*_y_*, σ*_c_* as standard deviation, respectively. The posterior distribution of σ*_y_* and σ*_c_* can be considered as a generalized description of the magnitude of multiplicative and additive platform effects. We can use them to sample the magnitudes of gene specific multiplicative and additive platform distortions for unobserved genes. The model is fitted for a given pair of scRNA-seq and targeted spatial transcriptomics platforms to account for the differences between them.

To check the extent to which the model is able to capture platform biases, we used three paired scRNA-seq and targeted spatial transcriptomics datasets: scRNA-seq and MERFISH data from mouse hypothalamic preoptic region [24] (Moffit dataset), scRNA-seq and osmFISH data from mouse cortex [26,59] (Codeluppi dataset), and scRNA-seq and DARTFISH data from healthy human kidney [60] (Zhang dataset) (**Methods,** **Table S1**). Fitting a model for each pair of datasets, we then performed posterior predictive check, i.e., we simulated spatial transcriptomics measurements from scRNA-seq data using the fitted Bayesian model (**Methods**). Comparisons of the distribution of simulated and observed spatial transcriptomics measurements demonstrated that the Bayesian model can accurately recapitulate the platform effects from different pairs of technologies (**Fig. 1E**, **Fig. S2A**–**C**). The posterior distributions of σ*_y_* and σ*_c_* (**Fig. 1F**–**G**) on the three datasets showed distinct levels of additive and multiplicative platform effects, indicating the need to account for platform-specific properties during gene panel selection.

### Gene panel selection using genetic algorithm

To take platform distortions into account during selection of the gene panels, we use the platform specific Bayesian model to simulate spatial transcriptomics measurements with distortions (**Methods**). The gene panels are optimized for their ability to recover cell type labels from such simulated spatial measurements, rather than the original scRNA-seq measurements. Such an approach is intended to provide a more accurate estimation of panel performance in a real spatially-resolved measurement. Instead of selecting top-performing genes, gpsFISH optimizes the entire gene panel in its combined ability to recover cell type labels. To optimize within this combinatorial gene space, gpsFISH uses genetic algorithm optimizer [61,62] (**Fig. 2**, **Methods**).

**Figure 2:**
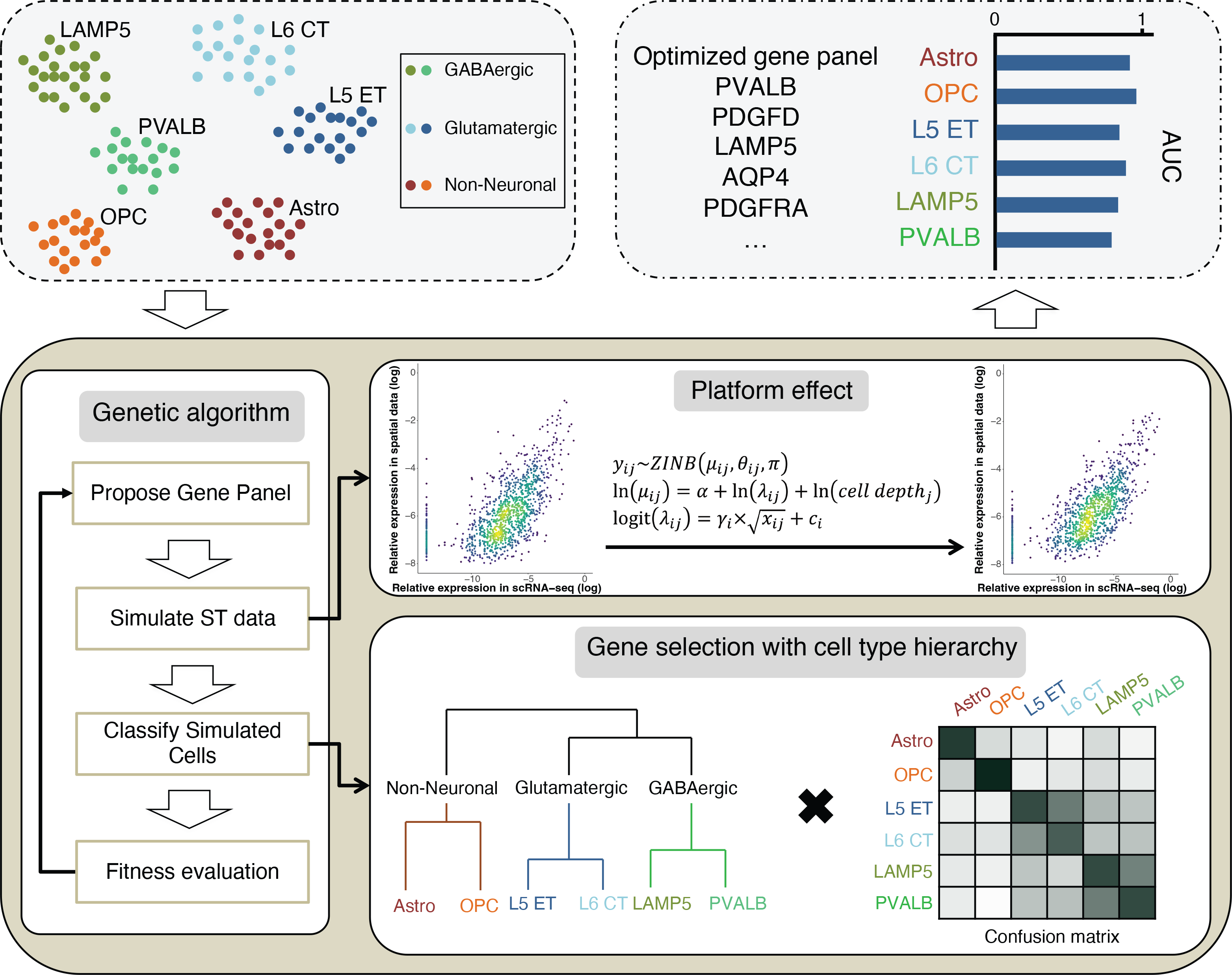
Schematic overview of gpsFISH. Upper left, an scRNA-seq dataset with cell type annotation is used as input. Bottom, a genetic algorithm framework is used for gene panel selection. Platform effects are accounted for using a Bayesian model. Cell type hierarchy can also be incorporated. Upper right, output includes optimized gene panel with classification statistics.

Within each iteration of optimization, multiple cross validations of classification are performed for each proposed gene panel. To avoid biasing towards a specific realization of spatial transcriptomics distortions, gpsFISH performs the platform simulations separately in each cross validation. As a result, gene panels that are more robust to unexpected platform distortions will be favored. This gene panel selection framework ensures the evaluation is reflective of the gene panel’s real classification performance when measured by specific targeted spatial transcriptomics technologies.

We first tested gpsFISH on the mouse hypothalamic scRNA-seq data (Moffitt dataset) with simulated platform effect by optimizing a 200 gene panel to distinguish “level 1” cell type annotation, which includes 12 broadly defined cell types (**Fig 3A**). Most of the cells are correctly classified, yielding an overall accuracy of 0.983 and high area under the receiver-operator curve (AUC) across all cell types (**Fig 3B**, **C**). The optimized gene panel selected with considering platform effect was also more successful in separating the 12 cell types on the resulting UMAP embedding compared to the gene panel selected without considering platform effect (**Fig. 3D**, **E**).

**Figure 3:**
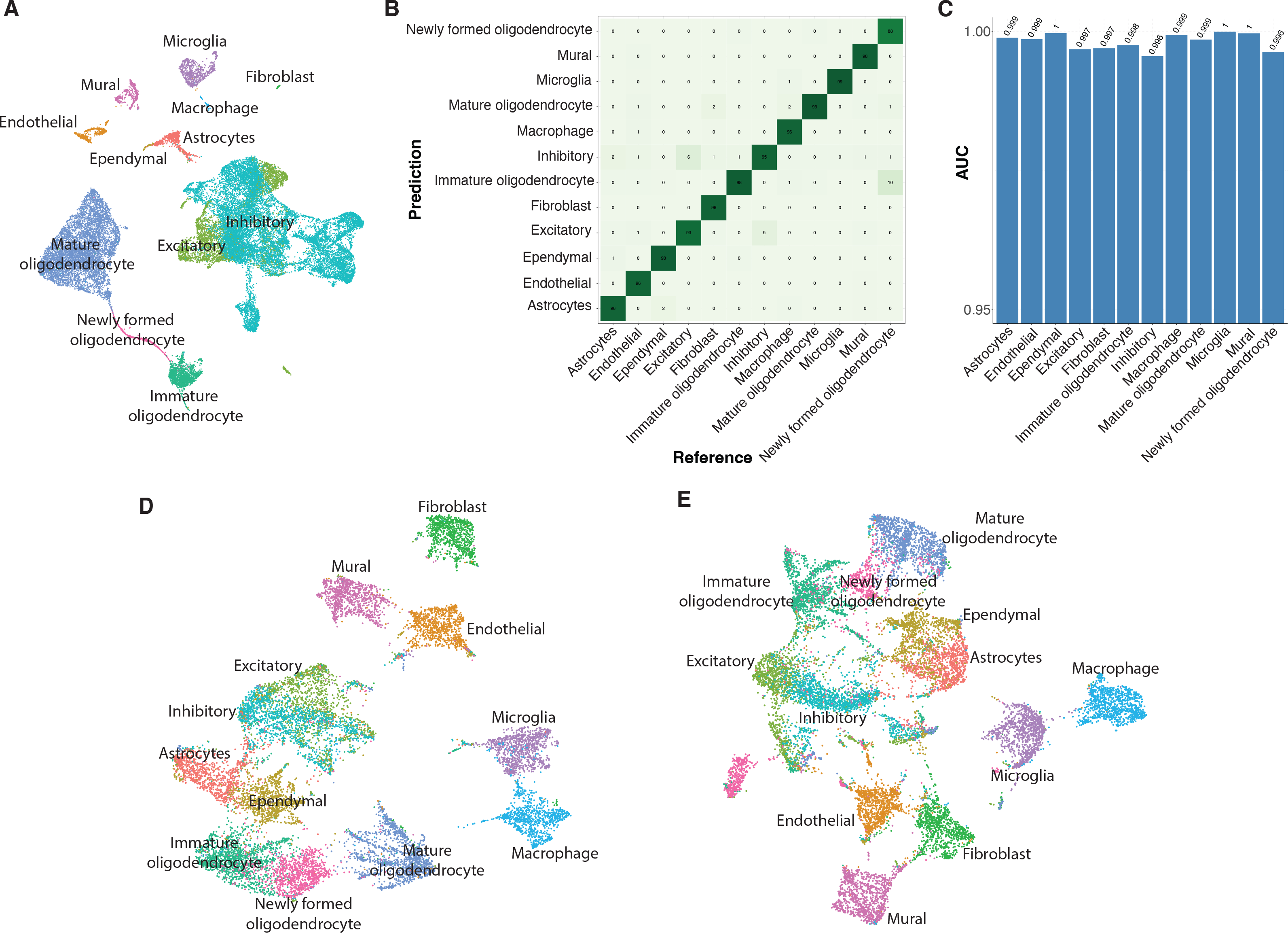
Gene panel selection using gpsFISH. **A**: UMAP of cells based on the mouse hypothalamic scRNA-seq data from Moffit dataset at level 1 cell type annotation. **B**: Normalized confusion matrix of the optimized gene panel for Moffit dataset at level 1 cell type annotation. **C**: AUC for each cell type of the same gene panel. **D-E**: UMAP of cells based on simulated spatial transcriptomics measurements with platform effect of the optimized gene panel selected with (**D**) and without (**E**)considering platform effect at level 1 cell type annotation.

To quantify performance of different methods we simulated spatial transcriptomics data from scRNA-seq, separating training and test sets (**Methods**). Simulations were performed both with and without distortions in order to evaluate how taking platform effects into account impacts gene panel performance. We also compared gpsFISH with two previously published classification-based gene selection methods: RankCorr and scGenefit. Both methods rely on the scRNA-seq expression profiles without considering platform effects. RankCorr is a rank-based one-vs-all feature selection method that selects marker genes for each cell type given a sparsity parameter, which controls the number of marker genes selected per cell type. We tuned this parameter to make sure the panels generated using RankCorr have the same size (200 genes). scGenefit selects gene markers that jointly optimize cell label recovery using label-aware compressive classification methods. As control, we provided a naïve way to simulate spatial transcriptomics measurements without platform effects (**Methods**). In addition, we also generated a panel of randomly selected genes as baseline.

The objective of gpsFISH optimization is to achieve high quality cell type classification on the spatial transcriptomics data. This entails two tasks: (1) selecting a good gene panel, and (2) using the gene panel for accurate cell type classification. In practice, while the design of an initial gene panel may rely on the scRNA-seq data, optimization of subsequent panels can take advantage of the probe-specific distortions that have already been observed in earlier measurements.

Similarly, as more and more spatial transcriptomics data are generated, when classifying cell types in a newly generated spatially-resolved measurement, it is likely that some partial annotations may already be available for that platform either on the current or previously acquired datasets. Regardless of the cell type granularity of partial annotation, it contains gene-specific platform effect information of genes in the spatial transcriptomics data, which can be estimated using our Bayesian model to improve cell type classification. Following this logic, we used two benchmark strategies, which evaluate the impact of platform effects on the two tasks (**Methods**). Both strategies share the same general framework in which a classifier is trained on the training data with gene expression profiles for all cell types, and then applied onto the testing data for cell type classification. The difference is how the two strategies incorporate partial annotation into the training data when available. Specifically, for the first strategy, evaluation with platform effect re-estimation (**Methods**), platform effects are estimated from the partial annotation data and incorporated into the training data for all gene selection methods. Since under this strategy the gene panels from all methods are evaluated in the same manner, it is useful in evaluating the impact of platform effect on the first task, i.e., selecting a good gene panel. In contrast, under the second evaluation strategy, evaluation without platform effect re-estimation (**Methods**), only gpsFISH panels are evaluated with platform effect estimation as described above (**Methods**), illustrating the impact of platform effects on both tasks.

Evaluation with platform effect re-estimation on the Moffit dataset using naïve Bayes as the classifier shows gpsFISH outperforms the control with naïve simulation and other gene selection methods (**Fig. 4A**), indicating that taking platform effects into consideration leads to more informative gene panels. Similar results were observed for the Zhang and Codeluppi dataset (**Fig. S3A** **and** **S3C**) and using random forest as classifier (**Fig. S4A**–**C**). From the normalized confusion matrix of the gene panel selected by gpsFISH with hierarchical tree on the left showing the relationship between cell types (**Fig. 4C**) we can see that most of the misclassifications are within the complex subpopulations of inhibitory and excitatory neurons.

**Figure 4:**
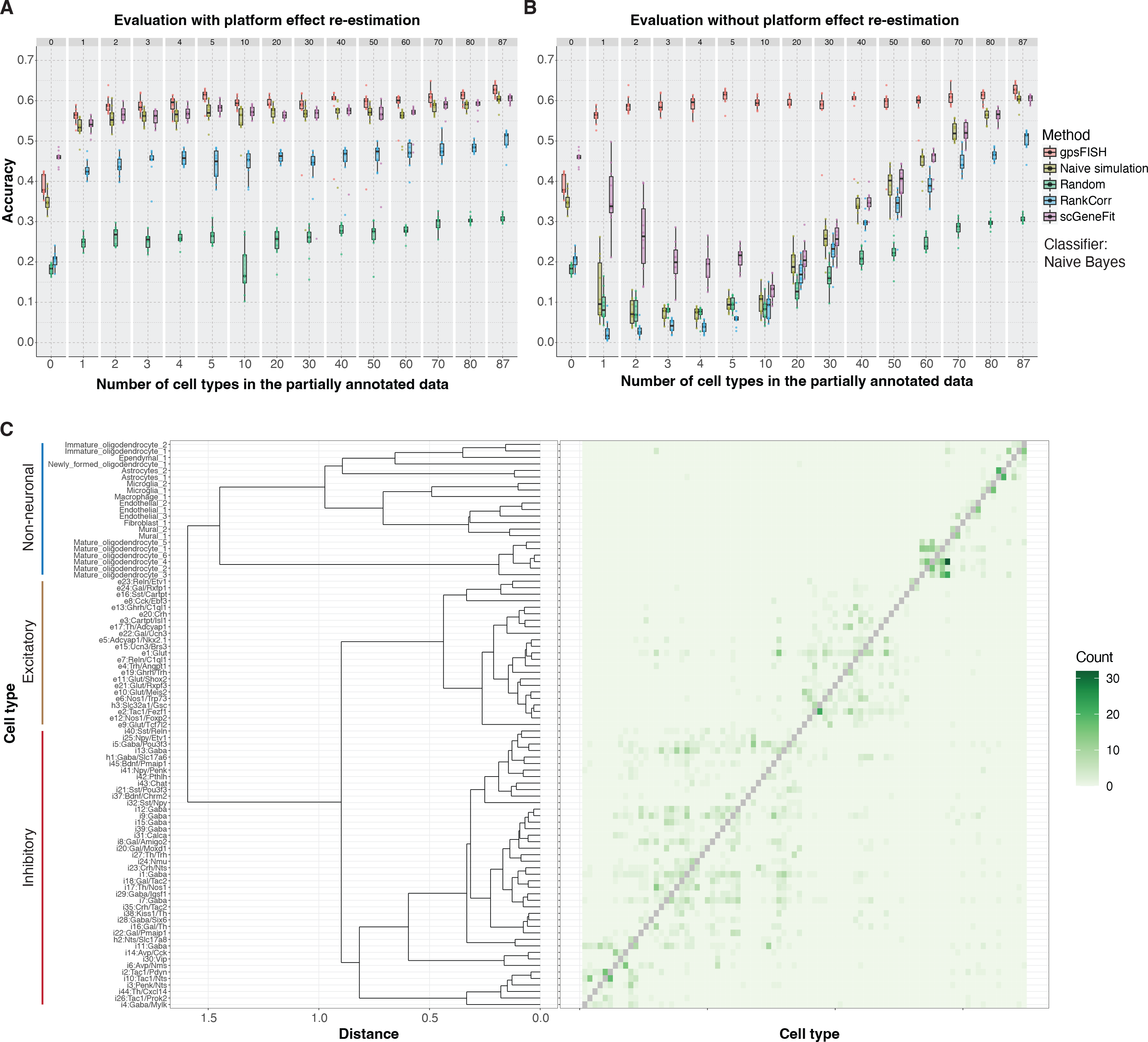
Comparison between gpsFISH and other gene selection methods. **A-B**: Box plot showing classification accuracy distribution of gene panels selected by 5 gene panel selection methods at different levels of partial annotation. The result is based on the Moffit dataset using evaluation with (**A**) and without (**B**) platform effect re-estimation. Naïve Bayes is used as classifier. **C**: Normalized confusion matrix of the optimized gene panel for Moffit dataset at level 2 cell type annotation with dendrogram showing the cell type hierarchy. Diagonal values of the confusion matrix are removed for better visualization of misclassifications.

A larger performance improvement of gpsFISH over other gene selection methods is observed using evaluation without platform effect re-estimation, especially when the level of partial annotation is low (**Fig. 4B**, **Fig. S3B** **and** **S3D**, **Fig. S4D**–**F**), indicating that considering platform effects can lead to more accurate cell type classification.

Overall, the comparison results show that gpsFISH outperforms other gene selection methods and considering platform effects can result in more informative gene panels and better cell type classification.

### Redundancy in gene space across independent gene panel optimizations enables incorporation of customized preferences

Independent panel optimizations performed multiple times (10) for each of the three datasets showed high level of redundancy in the gene space (**Fig. 5A**). Specifically, despite similar levels of overall performance, the overlap between independently optimized 200-gene panels was around 85, 65, and 35 genes, and more than 20%, 30%, and 45% of the genes showed up in only one of the 10 optimized gene panels for the Zhang, Moffit, and Codeluppi datasets, respectively (**Fig. S5A**–**C**). We observed similar level of redundancy even when the optimization was performed for a more granular “level 2” cell type annotations (46, 87, and 47 cell types for Zhang, Moffit, and Codeluppi dataset, **Fig. S5D**). The ability to achieve similar level of performance with different gene sets suggests that the panels can be further optimized to accommodate secondary criteria, such as inclusion of pre-selected genes, emphasis on genes with specific features or from specific pathways, etc.

**Figure 5:**
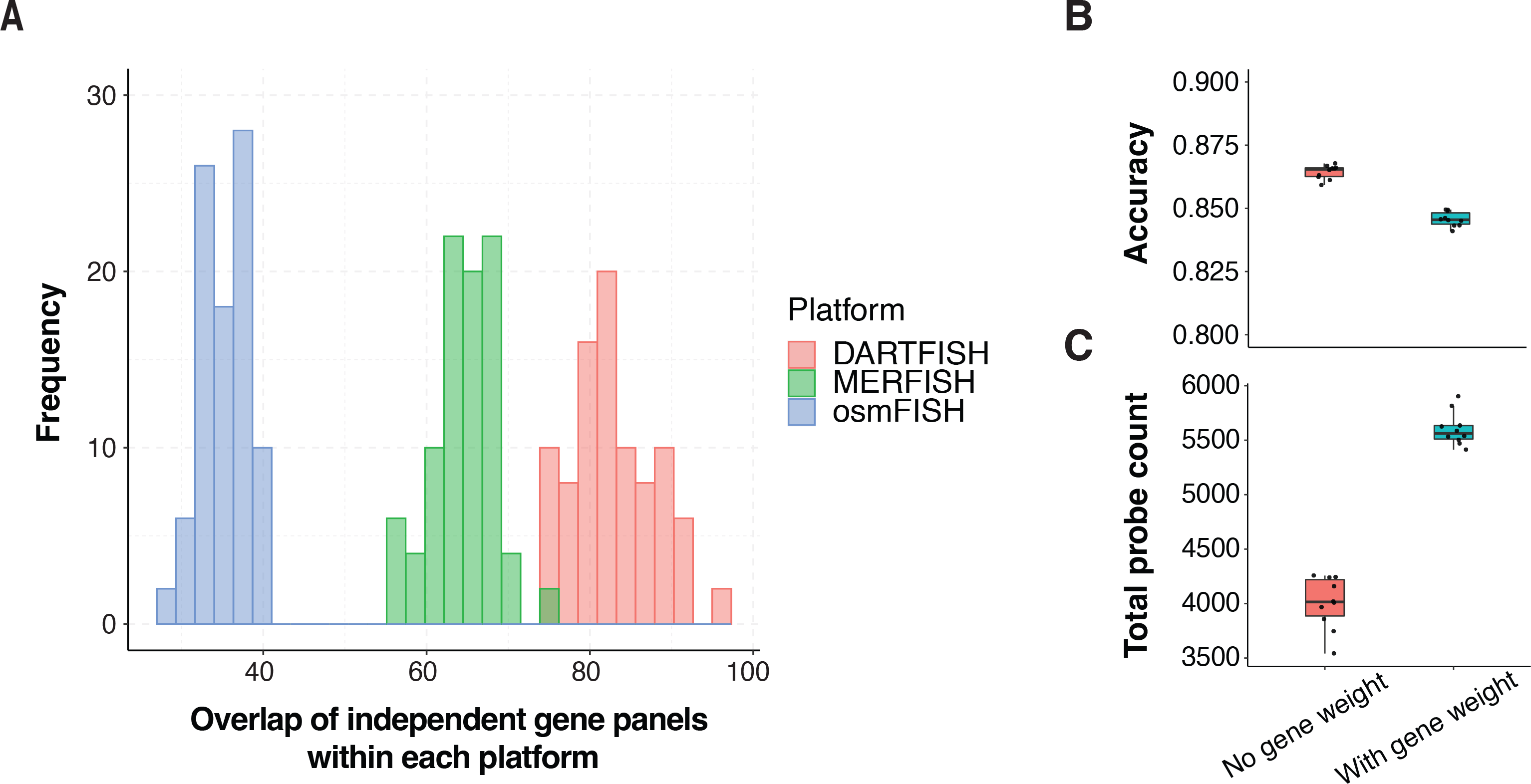
Redundancy in gene space across independent gene panel optimizations enables incorporation of customized preferences. **A**: Distribution of overlap of independent gene panels across 10 optimizations within each platform at level 1 cell type annotation. **B**: Accuracy of optimized gene panels without vs. with gene weight across 10 optimizations. **C**: Total number of probes of optimized gene panels without vs. with gene weight across 10 optimizations.

gpsFISH allows to incorporate secondary preferences during gene panel optimization, by specifying custom gene weights. To illustrate how panel redundancy can be used to incorporate secondary preferences with little impact on the classification performance, we evaluated the ability to increase the number of technical probes per gene. Specifically, many ISH-based assays, including DARTFISH, can include multiple different probes to enhance detection of any given transcript. The number of probes that can be designed to target each gene is determined by gene-specific factors like gene length. Genes with more probes are preferred, as they can be used to improve robustness and sensitivity of detection. To generate a gene panel with high number of potential probes, we used the predicted number of probes for each gene in the DARTFISH data (Zhang dataset) (**Methods**) as gene weight during gene panel selection. Of note, we capped the probe count at 15 to avoid bias towards a small portion of genes with extremely high number of probes (**Fig. S6A**). This also agrees with the fact that sensitivity will saturate when we have enough probes for a gene.

Following this approach, we performed 10 optimizations with and without probe count gene weights on the Zhang dataset using “level 1” cell type annotations. As expected, the optimizations with gene weight had slightly lower accuracy (**Fig. 5B**) but achieved a significantly higher number of total probes (**Fig. 5C**). This demonstrates that the redundancy of gene spaces allows one to incorporate additional customized constraints/preferences based on orthogonal information to design gene panels with preferred features without sacrificing the overall cell type classification performance.

### Hierarchical gene selection based on cell type hierarchy

Cell types are organized in a hierarchical manner with broad cell types divided into more detailed subpopulations. This hierarchical relationship can be considered when evaluating cell classification errors. For example, failure to distinguish two closely related subtypes, such as Th1 and Th17, of cells is likely to be considered less severe than mis-annotation of a Th cell into a different major cell type such as B cells.

In addition to the default “flat” cell type evaluation, gpsFISH therefore, implements a hierarchical classification option (**Fig. 6A**, **Methods**), in which correct classifications or misclassification between different cell types will receive different credit/penalty specified by a weighted penalty matrix according to cell type hierarchy. Using this hierarchical classification framework, gpsFISH provides flexibility to customize optimization based on desired level of cell type granularity.

**Figure 6:**
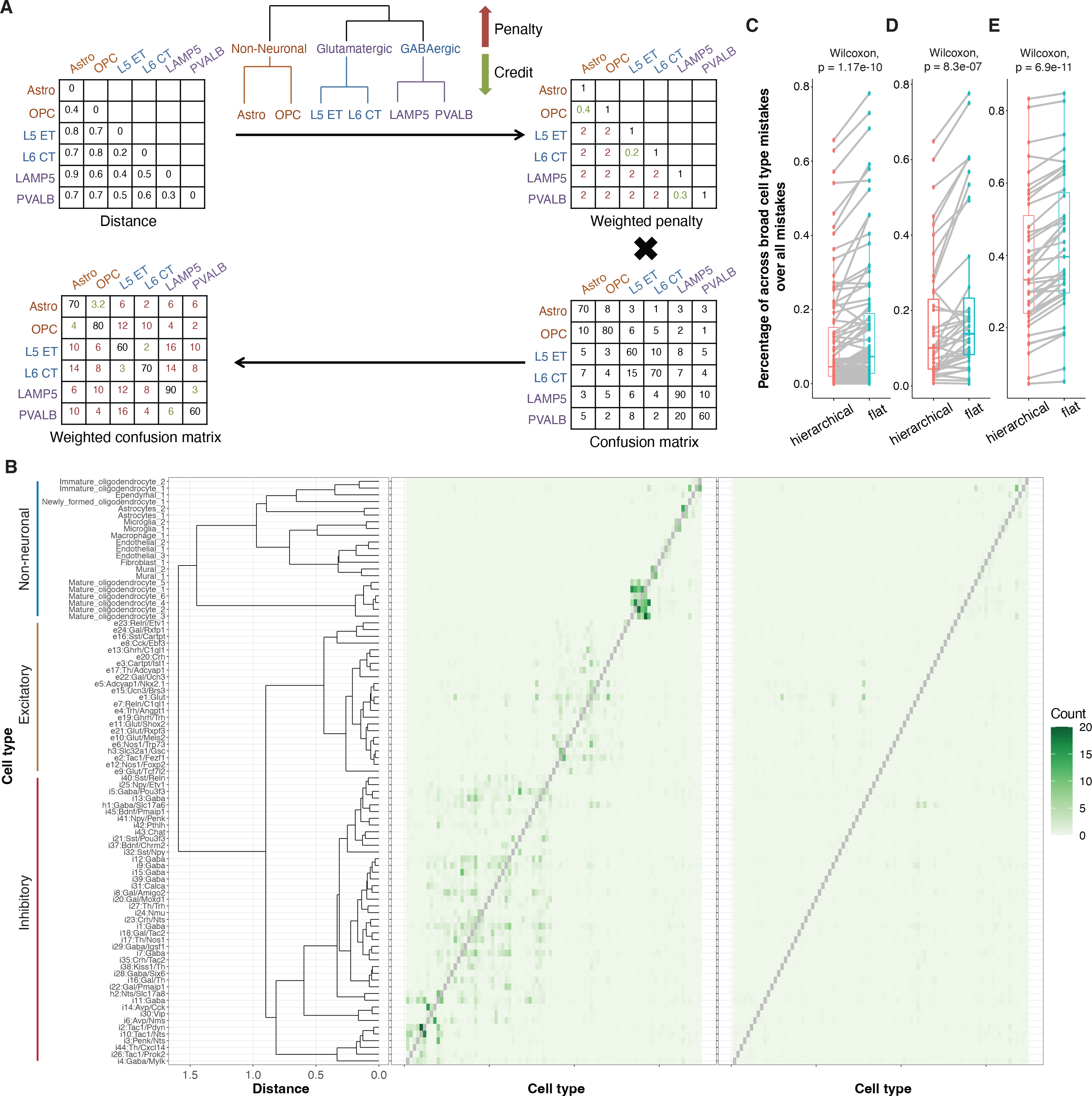
Gene panel selection with cell type hierarchy. **A**: Schematic of hierarchical gene selection using cell type hierarchy. A weighted penalty matrix is constructed using cell type hierarchy information quantified by pairwise distance between cell types. Additional penalty can be specified according to the cell type hierarchy. The weighted penalty matrix is then multiplied element-wise with the original confusion matrix to get the weighted confusion matrix for fitness evaluation. **B**: Original (left) vs. weighted (right) confusion matrix of the same optimized gene panel from Moffit dataset at level 2 cell type annotation with dendrogram showing the cell type hierarchy. Diagonal values of the confusion matrix are removed for better visualization of misclassifications. **C-E**: Percentage of across broad cell type (level 1) misclassifications over all misclassifications for flat vs. hierarchical classification on the Moffit (**C**), Codeluppi (**D**), and Zhang (**E**) dataset. Each dot represents one cell type with dots representing the same cell type connected. Wilcoxon paired test is performed between the percentages from flat vs. hierarchical classification and the p value is shown.

To evaluate the effect of the hierarchical classification for gene selection, we performed hierarchical gene selection at level 2 cell annotation of all three datasets. Under a hierarchical penalty scheme, misclassifications of cells between different level 1 categories incur a fixed penalty, whereas misclassifications within the same level 1 category were given partial credit, proportional to the expression similarity between the called and true subtypes (**Methods,** **Fig. 6B**). To quantify to what extent this hierarchical classification framework reduces misclassifications across broad cell types at level 1, we calculated the percentage of across broad cell type mistakes over all mistakes (**Methods**). We observed that the optimized gene panels using hierarchical classification tend to make significantly fewer misclassifications across broad cell types at level 1 compared to flat classification (**Fig. 6C**–**E**), indicating that cell type granularity can be controlled through the hierarchical classification framework.

## Discussion and conclusions

Accurate cell type classification is crucial for understanding the spatial relationship of cells in complex tissues. We implemented gpsFISH, a method for gene panel design of targeted spatial transcriptomics. By accounting for platform effects between scRNA-seq and targeted spatial transcriptomics technologies, gpsFISH is able to find more robust and informative gene panels and achieve better cell type classification.

Different technology has different patterns of platform effects. Specifically, we decomposed platform effects into two components: multiplicative and additive platform effects. While the multiplicative effect has been considered in deconvolution contexts (e.g., RCTD [47]), neither type of platform-specific distortions have been considered by other gene selection methods. Among other things, the additive platform effect enables gpsFISH to describe situations where specific genes show no expression in scRNA-seq data, but is detected in spatial transcriptomics data (dots forming the vertical line in **Fig. 1A** **and** **1B**). This observation is common for osmFISH (Codeluppi dataset) and MERFISH (Moffit dataset), and cannot be modelled using only multiplicative platform effect.

Comparing the three targeted spatial transcriptomics platforms, we found highest levels of additive platform effects in DARTFISH, followed by osmFISH and MERFISH (**Fig. 1G**). More specifically, DARTFISH had the lowest μ_#_, indicating the highest level of signal reduction compared to MERFISH and osmFISH (**Fig. S2E**). Signal reduction increases the possibility of good marker genes from scRNA-seq losing cell type specificity in spatial transcriptomics data (dots forming the horizontal line in **Fig. 1C**), which is a main scenario where platform effects affect gene panel selection. Higher level of signal reduction for DARTFISH agrees with our result that the performance improvement of gpsFISH over other gene selection methods is the largest in the Zhang dataset compared to the other two datasets, indicating the necessity to account for additive platform effects, especially for targeted spatial transcriptomics technologies with higher level of signal reduction.

In addition to additive platform effect, multiplicative platform effect also contributes to the systematic difference of transcripts detection rate across technologies, posing a challenge when transferring cell type information from scRNA-seq to spatial transcriptomics technologies. Comparison of three targeted spatial transcriptomics platforms shows osmFISH has the highest level of multiplicative platform effect, followed by MERFISH and then DARTFISH (**Fig. 1F**). Higher level of multiplicative platform effect leads to poorer cell type classification when there is no or low level of partial annotation compared to high level of partial annotation (**Fig. 4A** **and** **4B**, **Fig. S3** **and** **S4**), especially for evaluation without platform effect re-estimation due to distorted expression profiles between scRNA-seq and targeted spatial transcriptomics technologies. For evaluation with platform effect re-estimation, low level of partial annotation provided limited statistical power to accurately estimate gene specific platform effects, thus not able to increase the classification performance. This reduced performance is gone when we have more than one cell type included in the partial annotation, indicating partial annotation of a few cell types is enough to enhance cell type classification if multiplicative platform effects are accounted for.

Redundancy across independent optimizations allows incorporation of customized preferences into gene selection. However, gene weight needs to be carefully specified to ensure no sacrifice on overall gene panel performance. For the result in **Fig. 5B** and **5C**, we capped the number of probes for each gene at 15. For cutoffs lower than 15, gene weight difference between genes are small, leading to gene panels with similar performance but also similar total number of probes. However, for cutoffs higher than 15, the optimization will bias towards a small group of genes with high probe count, resulting in local minimum during optimization (**Fig. S6B**–**C**). This does achieve panels with significantly higher total number of probes, but the classification accuracy is dropped. This emphasizes the need to test different ways for gene weight specification in order to get the expected result without sacrificing performance.

Similarly, in our test of hierarchical gene selection, we specified the weighted penalty matrix directly from cell type hierarchy. Although we reduced misclassifications across broad cell types, the overall accuracy is slightly lower than flat classification (**Fig. S7**). This shows that partial credit of misclassifications needs to be given carefully, especially when there are many similar subpopulations within the same broad cell type like in the Moffit dataset. In real usage, it is suggested to prune the weighted penalty matrix constructed from the cell type hierarchy to remove unnecessary partial credit. Gene panel selection using flat classification can be run first to help adjust the weighted penalty matrix constructed using cell type hierarchy. In addition, the hierarchical classification provides a generic framework to fine tune emphasis of classification on certain cell types. Here we showed its usage to incorporate cell type hierarchy, but it is not restricted to cell type hierarchy. Customized weighted penalty matrix can be constructed using other information that provides preferences towards different classifications.

A major goal of spatial transcriptomics is to understand the spatial distribution of cell types and their corresponding cellular environment. gpsFISH facilitates this by selecting more informative and robust gene panels and providing ways for better cell type annotation. We also provide options to account for various custom preferences. As more targeted spatial transcriptomics data are generated, we expect that gpsFISH can facilitate the study of cellular organization of complex tissues under different biological contexts.

## Methods

### Datasets

In our study, we used three datasets that have both scRNA-seq and targeted spatial transcriptomics data from the same tissue. Information regarding the three datasets is summarized in **Table S1**. Further processing details are discussed below.

#### Moffit dataset

scRNA-seq data was downloaded from Gene Expression Omnibus (GEO) [63] under accession code GSE113576. MERFISH data was downloaded from Dryad [64]. Of note, the MERFISH data from Dryad is normalized and batch corrected. We undid the volume normalization and batch correction to get the original data.

In the scRNA-seq data, we first filtered out cells annotated as “Ambiguous” and “Unstable”. We then used information in the supplementary Table 1 of the original study to assign cell types. “Cell class (determined from clustering of all cells)” column was used as level 1 cell type annotation. “Neuronal cluster (determined from clustering of inhibitory or excitatory neurons)” and “Non-neuronal cluster (determined from clustering of all cells)” were used as level 2 cell annotation. Normalization was performed as described in the original study.

Only MERFISH data from naïve mice was used (to match scRNA-seq data). In addition, we also filtered out cells annotated as “Ambiguous” and “Unstable”. Fos gene and five blank genes were filtered out. 135 genes imaged in the combinatorial smFISH imaging were kept. Following the naming of cell types in Fig. 3D of the original study, we modified the cell type annotation of MERFISH data to make it consistent with the scRNA-seq data. Specifically, at level 1 cell type annotation, cells annotated as “Endothelial 1”, “Endothelial 2”, “Endothelial 3” were merged into “Endothelial”. “Astrocyte” was changed to “Astrocytes”. “OD Immature 1” and “OD Immature 2” were changed to “Immature_oligodendrocyte”. “OD Mature 1”, “OD Mature 2”, “OD Mature 3”, and “OD Mature 4” were changed to “Mature_oligodendrocyte”. “Pericytes” was changed to “Mural”. At cell type level 2, “Endothelial 1”, “Endothelial 2”, and “Endothelial 3” were changed to “Endothelial_1”, “Endothelial_2”, and “Endothelial_3”, respectively. “Ependymal” was changed to “Ependymal_1”. “OD Immature 1” and “OD Immature 2” were changed to “Immature_oligodendrocyte_1” and “Immature_oligodendrocyte_2”, respectively. “OD Mature 1”, “OD Mature 2”, “OD Mature 3”, and “OD Mature 4” were changed to “Mature_oligodendrocyte_1”, “Mature_oligodendrocyte_2”, “Mature_oligodendrocyte_3”, and “Mature_oligodendrocyte_4”, respectively.

After the processing above, additional filters were applied on the raw and normalized scRNA-seq data before gene panel selection. Genes with maximum cell type average expression lower than 1 were filtered out. In addition, long non-coding RNAs were also removed. As a result, 2886 and 5100 genes were used for gene panel selection at level 1 and 2, respectively. For platform effects estimation, the subset of the raw scRNA-seq and MERFISH data with cells from overlapping cell types were used.

#### Codeluppi dataset

scRNA-seq data was downloaded from GEO under accession code GSE60361. Annotation data was downloaded from [65]. osmFISH and corresponding annotation data was downloaded from [66].

For scRNA-seq data, cell labels in row 9 of the annotation were used as level 1 cell type annotation, and row 11 were used as level 2 cell type annotation. However, the level 1 cell type annotation is too broad (only 5 major cell types). Therefore, we regenerated level 1 cell type annotation by merging similar cell types at level 2 following descriptions from the original study. Specifically, in generating data for gene panel selection at level 1, “S1PyrDL”, “S1PyrL23”, “S1PyrL4”, “S1PyrL5”, “S1PyrL5a”, “S1PyrL6”, S1PyrL6b”, “ClauPyr” were merged into “S1_Excitatory”. “CA1Pyr1”, “CA1Pyr2”, “CA1PyrInt”, “CA2Pyr2”, “SubPyr” were merged into “Hippocampus_Excitatory”. 16 subclasses of interneurons (“Int1” to “Int16”) were merged into “Interneuron”. “Astro1” and “Astro2” were merged into “Astrocyte”. “Mgl1” and “Mgl2” were merged into “Microglia”. “Pvm1” and “Pvm2” were merged into “Pvm”. Six subpopulations of oligodendrocytes (“Oligo1” to “Oligo6”) were merged into “Oligodendorcyte”. “Vend1” and “Vend2” were merged into “Endothelial”. To make cell type labels consistent between scRNA-seq and osmFISH, “Peric” was changed to “Pericyte”. “Choroid” was changed to “Ventricle”. “Epend” was changed to “Ependymal”.

To generate the data for platform effect estimation, cell type labels were modified slightly differently to reflect the correspondence between cell types in scRNA-seq and osmFISH as shown in Fig. 2C and Fig. 2D of the original study. Specifically, three CA1 subclasses (“CA1Pyr1”, “CA1Pyr2”, “CA1PyrInt”) were merged into “Hippocampus_Excitatory”. 16 subclasses of interneurons (“Int1” to “Int16”) were merged into “Interneuron”. Two subclasses of microglia (“Mgl1” and “Mgl2”) were merged into “Microglia”. Two subclasses of perivascular macrophages (“Pvm1” and “Pvm2”) were merged into “Pvm”. Subclasses of S1 pyramidal cells were also merged: “S1PyrL4” and “S1PyrL5a” were merged into “S1_Excitatory_L45a”, “S1PyrL5” and “S1PyrL6b” were merged into “S1_Excitatory_L56b”. In addition, to make the cell type labels consistent between scRNA-seq and osmFISH, we changed “Astro1” and “Astro2” to “Astrocyte1” and “Astrocyte2”, respectively. We changed “Oligo6” to “Oligo_Mature”, “Oligo5” to “Oligo_MF”, “Oligo1”, to “Oligo_COP”, “Vend1” to “Endothelial1”, “Vend2” to “Endothelial2”, “Peric” to “Pericyte”, “Choroid” to “Ventricle”, “Epend” to “Ependymal”, “S1PyrL23” to “S1_Excitatory_L23”, and “S1PyrL6” to “S1_Excitatory_L6”. Cell types with fewer than 50 cells were removed.

For osmFISH data, we first filtered out invalid cells based on the “Valid” column of the annotation data. Then, similar to scRNA-seq data, we modified cell type labels according to Fig. 2C and Fig.2D in the original study, which shows correspondence between cell types in scRNA-seq and osmFISH. Specifically, “Astrocyte Gfap” was changed to “Astrocyte1”. “Astrocyte Mfge8” was changed to “Astrocyte2”. “Hippocampus” was changed to “Hippocampus_Excitatory”. “pyramidal L4” was changed to “S1_Excitatory_L45a”. “Pyramidal L5” was changed to “S1_Excitatory_L56b”. “Pyramidal L6” was changed to “S1_Excitatory_L6”. “Perivascular Macrophages” was changed to “Pvm”. “Oligodendrocyte COP” was changed to “Oligo_COP”. “”Oligodendrocyte Mature”” was changed to “Oligo_Mature”. “Oligodendrocyte MF” was changed to “Oligo_MF”. “Endothelial 1” was changed to “Endothelial1”, and “Endothelial” was changed to “Endothelial2”. “Pericytes” was changed to “Pericyte”. “Vascular Smooth Muscle” was changed to “Vsmc”, “C. Plexus” was changed to “Ventricle”. “Pyramidal L2-3” and “Pyramidal L2-3 L5” were merged into “S1_Excitatory_L23”. “Inhibitory Cnr1”, “Inhibitory CP”, “Inhibitory Crhbp”, “Inhibitory IC”, “Inhibitory Kcnip2”, “Inhibitory Pthlh”, and “Inhibitory Vip” were merged into “Interneuron”.

scRNA-seq data was normalized using the count_normalize function in the scran package. Similar to the Moffit dataset, the raw and normalized scRNA-seq were further filtered before gene panel selection using the same filters. 6123 and 9052 genes were used for gene panel selection at level 1 and 2, respectively. For platform effect estimation, the subset of the raw scRNA-seq and osmFISH data with cells from overlapping cell types were used.

#### Zhang dataset

Raw and normalized scRNA-seq data from kidney were obtained from [60]. They were further filtered before gene panel selection using the same filters. 2920 and 3796 genes were used for gene panel selection at level 1 and 2, respectively. The DARTFISH data is unpublished. It can be found in Zenodo [67]. We annotated the cells in the DARTFISH data manually using curated marker genes (**Table S2**) at subclass level (third column). For platform effect estimation, the subset of the raw scRNA-seq and DARTFISH data with cells from overlapping cell types were used.

### Platform effects estimation using a Bayesian model

We assume the observed number of molecules *y_ij_* in the spatial transcriptomics data for gene *i* in cell *j* follows a zero-inflated negative bimonial (ZINB) distribution with:

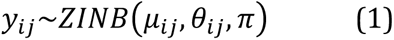

where π is the zero inflation parameter which is assumed to be constant across genes and cells. μ*_ij_* is the mean parameter determined by a global intercept α, true expression level of gene *i* in cell *j* denoted as λ*_ij_*, and the cell depth (total number of molecules) of cell *j* from spatial transcriptomics data as 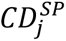:

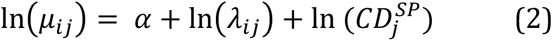

To account for platform effects, we assume the true expression level λ_!$_ is a random variable defined by:

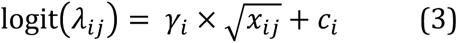

where γ*_i_* is a gene specific coefficient representing multiplicative platform effects, and *C_i_* is a gene specific intercept representing additive platform effects. *x_ij_* represents the relative expression of gene *i* in cell *j* calculated from scRNA-seq data:

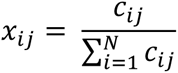

where *C_ij_* is the number of count for gene *i* in cell *j* from the scRNA-seq data, and *N* is the totol number of genes. When fitting the Bayesian model, in order to match measurement between scRNA-seq data and targeted spatial transcriptomics data, we used cell type average relative expression to replace individual cell level relative expression:

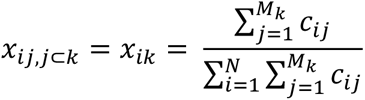

where *M_k_* is the number of cells in cell type *k*.

For the dispersion parameter θ*_ij_* of the ZINB distribution, we assume it is also dependent on λ*_ij_*:

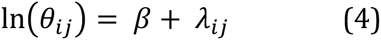

where β is the intercept.

A Beta prior distribution is assumed for π. For α, β, and *C_i_*, we assume they follow normal distribution. γ*_i_* is assumed to follow a log-normal distribution:

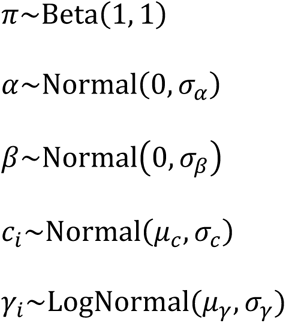

where the hyperparameters are assumed to follow Cauchy and half Cauchy distribution:

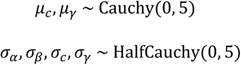

scRNA-seq and targeted spatial transcriptomics data from overlapping genes and overlapping cell types were used as input. Additional filters were applied on the MERFISH data to reduce the totol number of cells for more efficient estimation. Specifically, cells with cell depth lower than 100 were filtered out. Cell types with fewer than 1000 cells were filtered out. Then we subsampled each cell type to keep at most 1000 cells for each cell type. Variational inference in Stan was used for model fitting.

### Simulation of spatial transcriptomics measurements from scRNA-seq data with platform effects

We used fitted Bayesian models to simulate spatial transcriptomics measurements from scRNA-seq data. Specifically, α, β, π, μ*_c_*, σ*_c_*, μ_γ_, σ_γ_ were randomly sampled from their estimated posterior distribution. *C_i_*, and γ*_i_* were randomly sampled from their corresponding normal and log normal distribution for each new gene that is not observed in the data used to fit the Bayesian model. If a gene is already seen during fitting the Bayesian model, we can either use the empirical *C_i_*, and γ*_j_* estimated during model fitting (used in this study) or randomly sample them from the corresponding normal and log norml distribution. 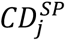 was randomly sampled from empirical cell depth distribution from observed targeted spatial transcriptomics data. *x_ij_* was calculated from scRNA-seq data. It can be cell type average as we used in model fitting or calculated within each individual cell. In our study, the latter was used when simulating spatial transcriptomics measurements to maintain the cell level heterogenity in scRNA-seq data. Finally, the generated values were plugged into equations (1), (2), (3), and (4) to generate spatial transcriptomics measurements.

### Simulation of spatial transcriptomics measurements from scRNA-seq data without platform effects (naïve simulation)

We provided a naïve way to simulate spatial transcriptomics measurements without platform effects as control. During the simulation without platform effects, cell depth of simulated spatial transcriptomics cell were randomly sampled from the empirical cell depth distribution of observed targeted spatial transcriptomics data. Of note, the empirical cell depth distribution was adjusted proportionally based on the ratio between relative expression of new genes for simulation and relative expression of overlapping genes used in fitting the Bayesian model. After having the simulated cell depth for each cell, the number of molecules for each gene within each cell was sampled from a multinomial distribution with size equal to the simulated cell depth and probability equal to each gene’s relative expression in that cell. At the end, genes were randomly selected given the probe failure rate. Then, simulated molecule count of selected genes were set to 0 to reflect probe failure.

### Genetic algorithm for gene panel selection

We used genetic algorithm as the framework for gene panel selection. Each individual in a population is one candidate gene panel. We set the gene panel size to 200 genes. Each population contains 200 individuals.

The first step of genetic algorithm is to initialize a population of candidate gene panels. The genes can be either randomly selected from all candidate genes or selected based on their differential expression between cell types. In this study, we took a hybrid approach. 95% of the 200 gene panels were initiated randomly from all candidate genes to maintain population diversity. The rest 5% were initialized using DEGs for each cell type. DE analysis was performed using Pagoda2. Genes with AUC greater than 0.7 were considered significant.

The second step is to evaluate the fitness of each candidate gene panel in the population. Here we define fitness as the average classification accuracy over 5 cross validations. Classification was performed on simulated spatial transcriptomics measurements from scRNA-seq data. Cell type annotation from scRNA-seq data was used as ground truth. The accuracy was calculated based on the original confusion matrix for flat classification and weighted confusion matrix for hierarchical classification. We provided two classifiers, random forest and naïve Bayes. In this study we used naïve Bayes due to its fast speed and relatively similar level of accuracy compared to random forest. To improve the efficiency, scRNA-seq data was subsampled to reduce the number of cells for large cell types and resampled to increase the number of cells for small cell types. Specifically, for level 1 cell type annotation, cell type size was capped at 1500 cells. The lower bound was set as 1000 cells for Moffit dataset and 500 for Zhang and Codeluppi dataset. For level 2 cell type annotation, 250 and 500 were used as the cell type size range for Moffit dataset. The range for Zhang and Codeluppi dataset was 300 and 900.

The third step is selection and mutation. The selection strategy we used is tournaments. Specifically, randomly selected candidate gene panels face each other 1 vs. 1. The one with a higher fitness value was used as parent. In addition, candidate gene panels with higher fitness values were more likely to be selected in the tournaments. After having the parent gene panels, uniform crossover was performed to generate the offspring gene panels. Duplicated genes after uniform crossover were replaced by randomly sampled genes in the parent candidate gene panels but not in the offspring gene panel. Mutation was then performed to maintain gene diversity and prevent premature convergence. We set the mutation rate to 1%. When gene weight was provided, genes with higher weight were (1) more likely to be selected during crossover, (2) less likely to be mutated if it is already in the population, (3) and more likely to be introduced into the population through mutation if it is not in the current population.

Finally, the same process was repeated for the offspring population. The candidate gene panel with the highest fitness value for one iteration was considered as the optimal gene panel. If the iteration after it has a candidate gene panel with higher fitness value, the optimal panel will be replaced by this new candidate gene panel. Otherwise, the optimal panel will stay the same. The iterative process will end either when it reaches a given number of iterations, or the accuracy doesn’t improve more than a threshold for a given number of iterations. In our study, we ran all the optimizations for at least 500 iterations to ensure convergence although in all cases the optimization converged a lot earlier.

If a list of pre-selected genes, e.g., canonical marker genes based on previous knowledge, is provided, genes in the list will be included in each candidate gene panel as well as the final optimal gene panel.

### Hierarchical classification using cell type hierarchy

During genetic algorithm optimization, a weighted penalty matrix can be provided to assign partial credit or extra penalty to classification between certain cell types. The weighted penalty matrix is a square matrix with each row and each column representing one cell type. For each value ρ*_ij_* (*i* ≠ *j*) in the weighted penalty matrix, if ρ*_ij_* > 1, an extra penalty is given to misclassifying cells from cell type *j* to cell type *i*. If ρ*_ij_* < 1, a partial credit is given to misclassifying cells from cell type *j* to cell type *i*. ρ*_ij_* = 1 means no penalty or partial credit. In hierarchical classification, the weighted penalty matrix was incorporated to the confusion matrix by element-wise multiplication to provide a weighted confusion matrix. The accuracy of the weighted confusion matrix was used to evaluate the fitness of candidate gene panels.

Essentially, the weighted penalty matrix can be constructed arbitrarily by user’s preference. In this study, we constructed the weighted penalty matrix from cell type hierarchy. First, pairwise distance between cell types was calculated. Specifically, average expression profile of each cell type was calculated using normalized count by taking average expression of all cells in each cell type. Top 1000 genes with highest standard deviation were used to calculate pairwise Pearson correlation coefficient. One minus the pairwise Pearson correlation coefficient was used as pairwise distance between cell types. Second, the pairwise distance matrix was normalized by the largest distance so the values range from 0 to 1. Third, the pairwise distance matrix was then adjusted based on cell type hierarchy.

Specifically, a level of cell type annotation was selected as reference. For cell types below the reference level that are from the same cell type at the reference level, the pairwise distance (between 0 and 1) between them was kept unchanged to reflect partial credit to wrong classifications among them. For cell types below the reference level that are from different cell types at the reference level, the pairwise distance between them was changed to a user defined value where 1 means no extra penalty and greater than 1 means extra penalty. In this study, we used 1 for no extra penalty and level 1 cell type annotation was used as reference. Finally, the diagonal value was changed to 1 to reflect no extra credit to correct classifications. This weighted penalty matrix was used for hierarchical classification in our study.

### Calculating the percentage of across broad cell type mistakes over all mistakes

We performed 5 optimizations with flat classification and hierarchical classification for all three datasets, respectively. Average confusion matrix over 5 optimizations for each data was calculated. After that, for each cell type, we counted the total number of misclassifications and among all the misclassifications, what proportion of them misclassifies cells to cell types at level 2 that don’t belong to the same cell type at level 1.

### Gene panel selection using RankCorr and scGeneFit

The same scRNA-seq data from the three datasets after filtering were used as input. For RankCorr, raw scRNA-seq data before normalization was used as suggested. The lamb parameter was tuned to make sure the output marker gene list has 200 genes. For scGeneFit, normalized scRNA-seq data was used by following the examples on its GitHub page. Panel size was set to 200.

### Evaluation of optimized gene panel

To evaluate optimized gene panels, we first simulated spatial transcriptomics measurements with platform effects based on the gene panel’s expression profile in scRNA-seq data. Then this simulated spatial transcriptomics data was split into training and testing data. The training data contains cells from a subset of cell types whose cell type labels are known. This was used as the partial annotation of the simulated spatial transcriptomics data. The testing data contains cells from all cell types (excluding cells in the training data), which is considered as part of the simulated spatial transcriptomics data that hasn’t been annotated yet. We varied the number of cell types in the training data from zero to all the cell types to reflect different levels of partial annotation. When there was no partial annotation, scRNA-seq data was used as the final training data for classifier training. When there was partial annotation, information in the partial annotation was included in the final training data. After that, a classifier (naïve Bayes or random forest) was trained using the final training data and applied on the testing data for cell type classification evaluation. Since the testing data was simulated from scRNA-seq data, the cell type labels in scRNA-seq data were used as ground truth. Classification accuracy was used as the metric to evaluate a gene panel. At each level of partial annotation, we repeated the same calculation 10 times. To separately evaluate the impact of platform effects on gene panel selection and cell type classification, within the same framework described here, we designed two different strategies to evaluate a gene panel by varying whether platform effect distortions that can be learned from partial annotation examples are used to produce more realistic training data for cell type classification.

#### Evaluation with platform effect re-estimation

This evaluation strategy was designed to focus on the performance of the optimized gene panels, and not on the differences in the cell type classification (evaluation) stage. In this strategy, partial annotation was first used to estimate gene specific platform effects using the Bayesian model. We then used these estimated gene specific platform effects to simulate an updated spatial transcriptomics training data, which will be combined with the partially annotated spatial transcriptomics data and then used for training cell type classifiers for all the methods being evaluated. Only cell types not already available in the partially annotated spatial transcriptomics data were simulated. When partial annotation was available for 5 or fewer cell types, the final training data combined the partially annotated and simulated spatial transcriptomics training data with scRNA-seq data. When more than 5 cell types were available, training was performed on the partially annotated and simulated spatial transcriptomics training data only. The final training data and testing data were normalized by the total number of transcripts within each cell and scaled by 10000. It was then log transformed after adding 1 pseudocount. This normalized training and testing data were used for classifier training and testing.

#### Evaluation without platform effect re-estimation

In this evaluation strategy, only gpsFISH is able to make use of the platform effects information in the partial annotation (as described above). All the other methods used the partial annotation according to their own method design. Specifically, for the control which used naïve simulation during gene panel selection, the empirical cell depth distribution of the complete testing data was used to simulate a spatial transcriptomics training data without platform effect. This simulated spatial transcriptomics training data was used in the same way as described above to get the final training data. For RankCorr, scGeneFit, and the random panel, since the gene selection was solely based on scRNA-seq data, cells in the partial annotation were directly combined with the scRNA-seq data of cell types not already available in the partial annotation. The combined data were used as the final training data. Same normalization was performed on the final training data and testing data before classifier training and testing.

### Calculate the number of probes for each gene for the DARTFISH data

During the generation of the DARTFISH data, ppDesigner [68] was used to calculate the number of probes that can be designed to target each gene.

## Declarations

### Ethics approval and consent to participate

Not applicable.

### Consent for publication

Not applicable.

### Availability of data and materials

Scripts to generate data and to perform the above analysis are available in Zenodo [67].

gpsFISH’s open-source code is maintained and documented on Github [69] and is publicly available under the MIT license.

Pre-fitted Bayesian models based on the Zhang, Moffit, and Codeluppi dataset respectively are deposited in Zenodo [70].

### Competing interests

P.V.K. serves on the Scientific Advisory Board to Celsius Therapeutics Inc. and Biomage Inc. P.V.K. is an employee of Altos Labs.

### Funding

P.V.K. and Y.Z. were supported by 5U54HL145608 grant from NIH.

### Authors’ contributions

P.V.K. and Y.Z. formulated the study and the overall approach. Y.Z. developed the detailed algorithms and performed the analysis with advice from P.V.K. and V.P. Y.Z. implemented the gpsFISH package with help from E.B. R.Q. and K.Z. generated the DARTFISH dataset. Y.Z. and P.V.K. drafted the manuscript.

## Acknowledgements

We thank Hirak Sarkar and Teng Gao for useful discussions on the method development. We also thank Chienju Chen and Kian Kalhor for useful discussions on the DARTFISH technology and data processing. We are also grateful to Jeffrey Moffitt who provided help with obtaining the original MERFISH data before normalization and batch correction. P.V.K. serves on the Scientific Advisory Board to Celsius Therapeutics Inc. and Biomage Inc. P.V.K. is an employee of Altos Labs.

## Supplementary Information

Additional file 1: Table S1.xlsx

Information of the Moffit, Codeluppi, and Zhang dataset

Additional file 2: Table S2.xlsx

Curated marker genes for the Zhang dataset

---

**Figure S1:**
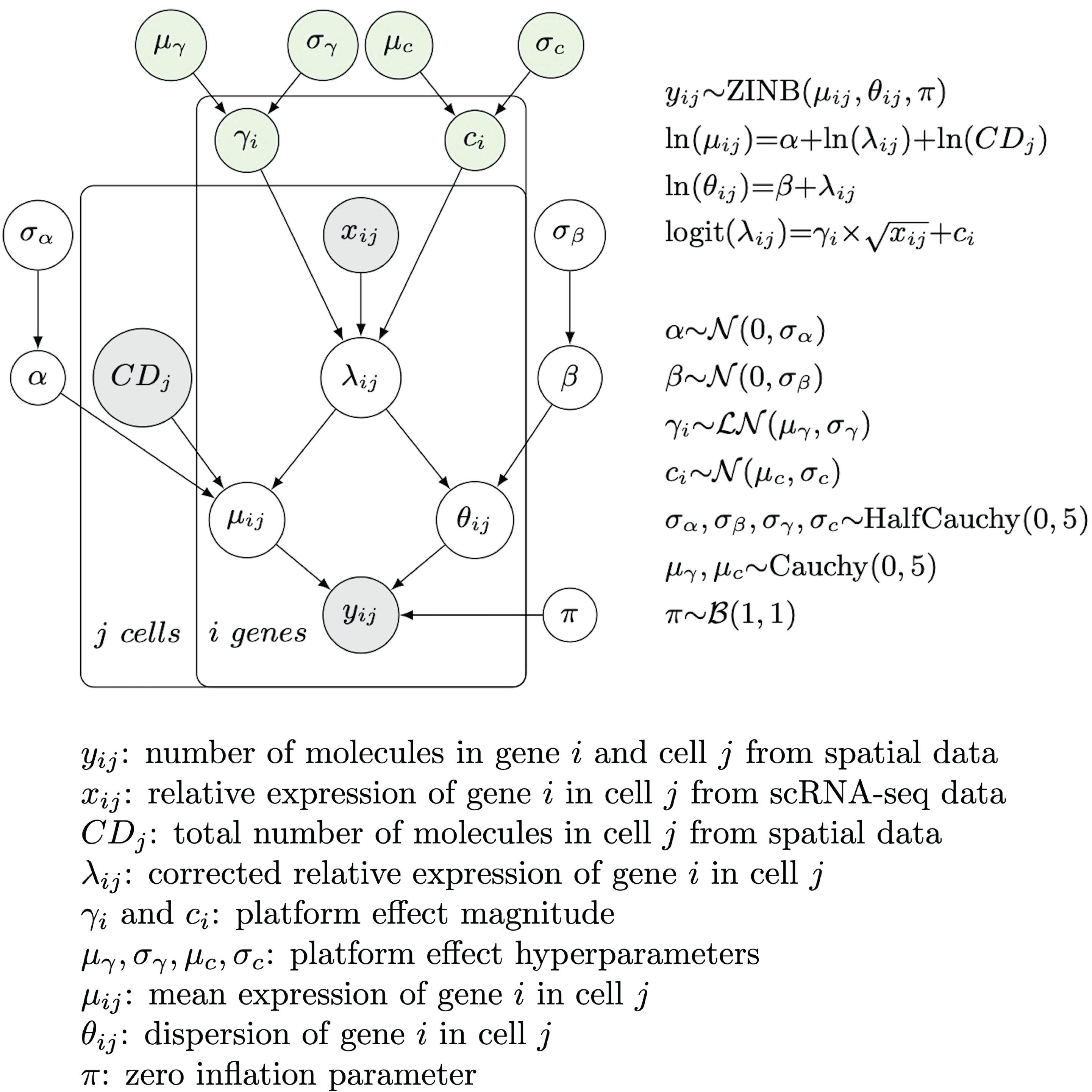
Schematic of the Bayesian model for platform effect estimation. Circles in gray represent observed variables. Circles in green correspond to platform effect related variables to be estimated.

**Figure S2:**
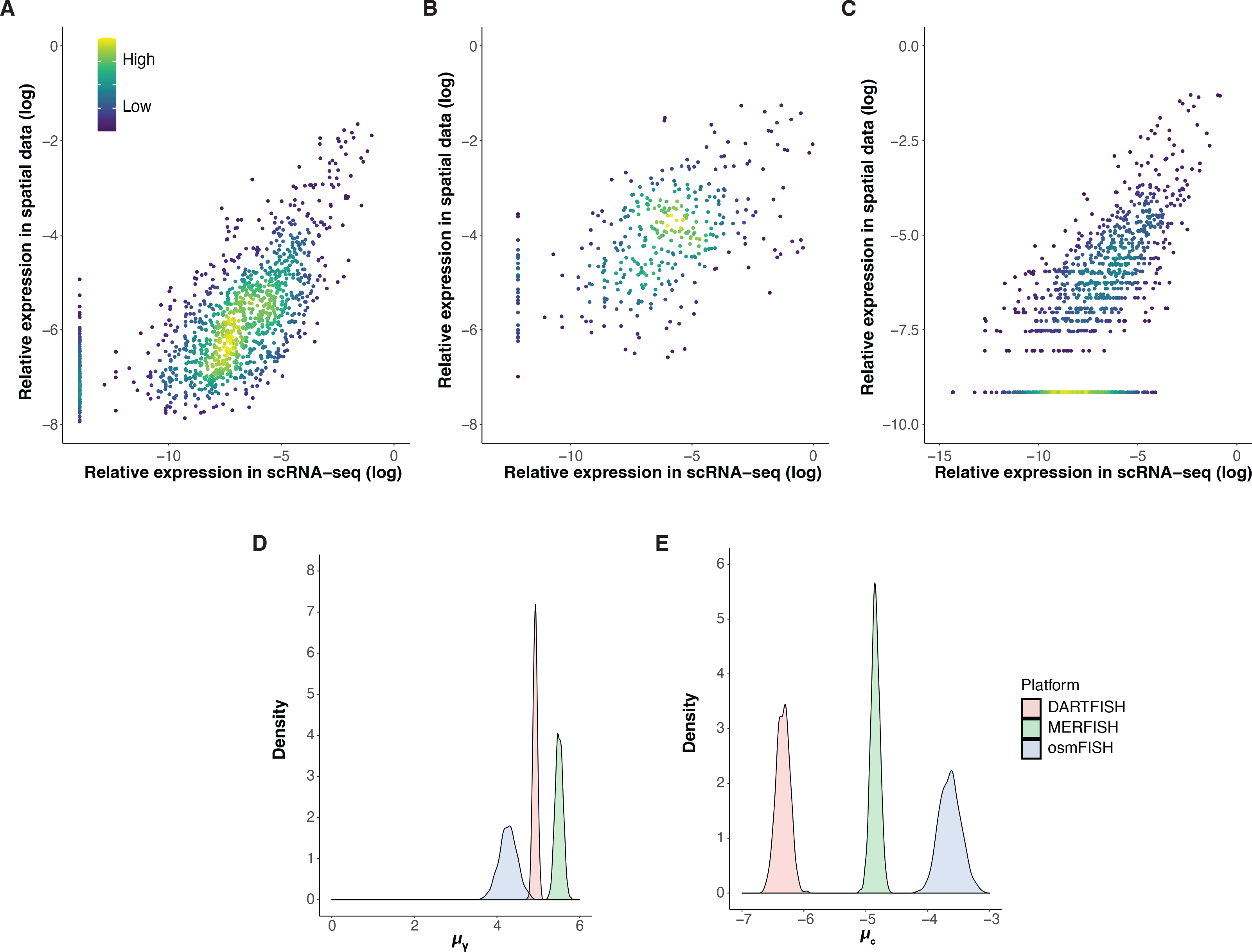
Bayesian model captures platform effect between scRNA-seq and targeted spatial transcriptomics technologies. **A-C**: Scatter plot showing the log transformed relative expression of genes measured by scRNA-seq vs. simulated spatial transcriptomics data using fitted Bayesian model for Moffit (**A**), Codeluppi (**B**), and Zhang (**C**), respectively. **D-E**: Density plot showing the estimated posterior distribution of μ_!_ (**D**) and μ_”_. (**E**).

**Figure S3:**
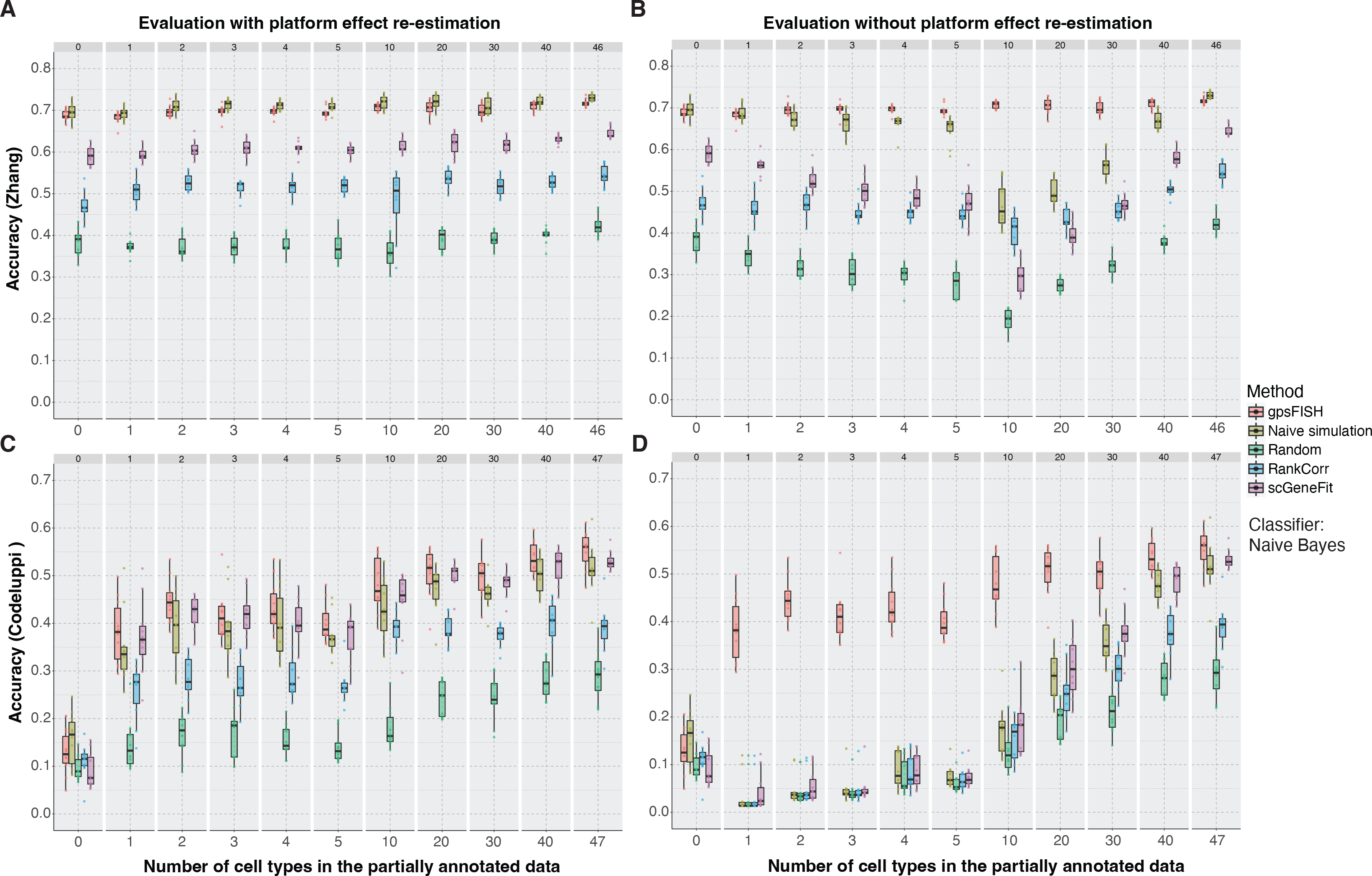
Comparison between gpsFISH and other gene selection methods on the Zhang and Codeluppi dataset. Box plot showing classification accuracy distribution of gene panels selected by 5 gene panel selection methods at different levels of partial annotation. (**A**) Zhang dataset using evaluation with platform effect re-estimation. (**B**) Zhang dataset using evaluation without platform effect re-estimation. (**C**) Codeluppi dataset using evaluation with platform effect re-estimation. (**D**) Codeluppi dataset using evaluation without platform effect re-estimation. Naïve Bayes is used as classifier.

**Figure S4:**
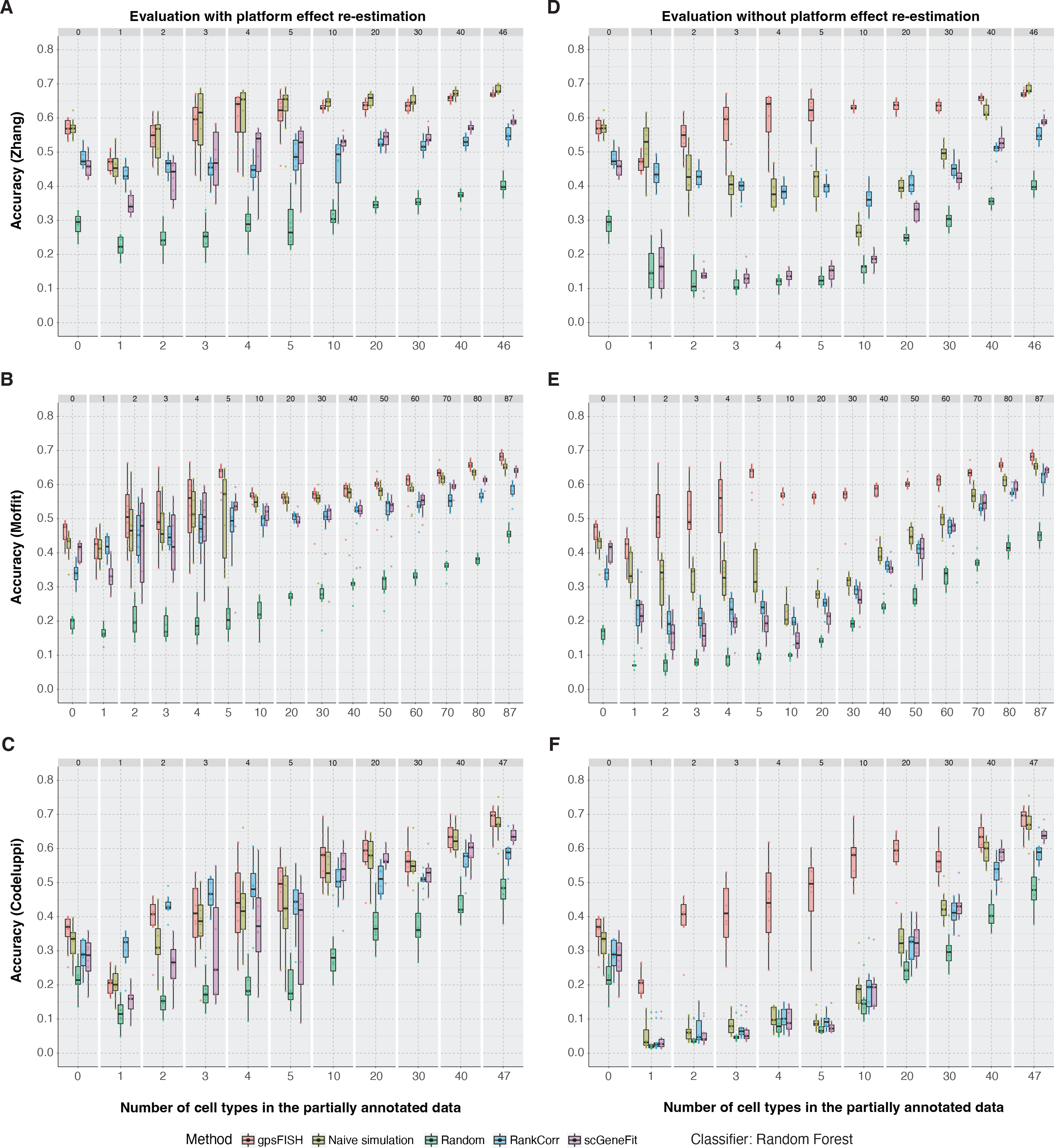
Comparison between gpsFISH and other gene selection methods using random forest as classifier. Box plot showing classification accuracy distribution of gene panels selected by 5 gene panel selection methods at different levels of partial annotation for the three datasets using evaluation with (**A-C**) and without (**D-E**) platform effect re-estimation. Random forest is used as classifier.

**Figure S5:**
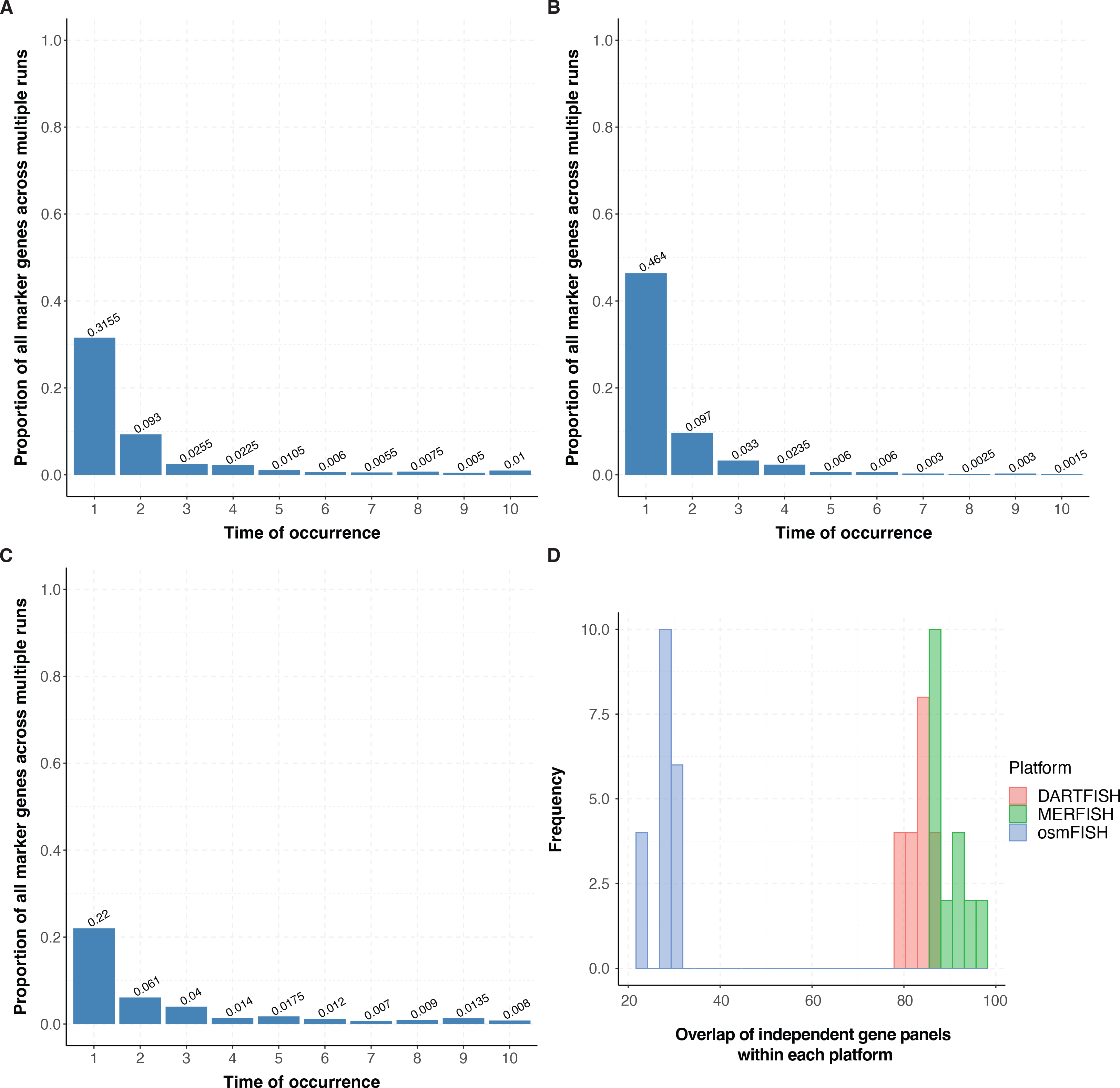
High redundancy across optimizations using gpsFISH. **A-C**: Bar plot showing among all the genes selected in 10 optimizations, the percentage of them that are included in 1 to 10 optimized panels for Moffit (**A**), Codeluppi (**B**), and Zhang (**C**) dataset at level 1 cell type annotation. **D**: Distribution of overlap of independent gene panels across 10 optimizations within each platform at level 2 cell type annotation.

**Figure S6:**
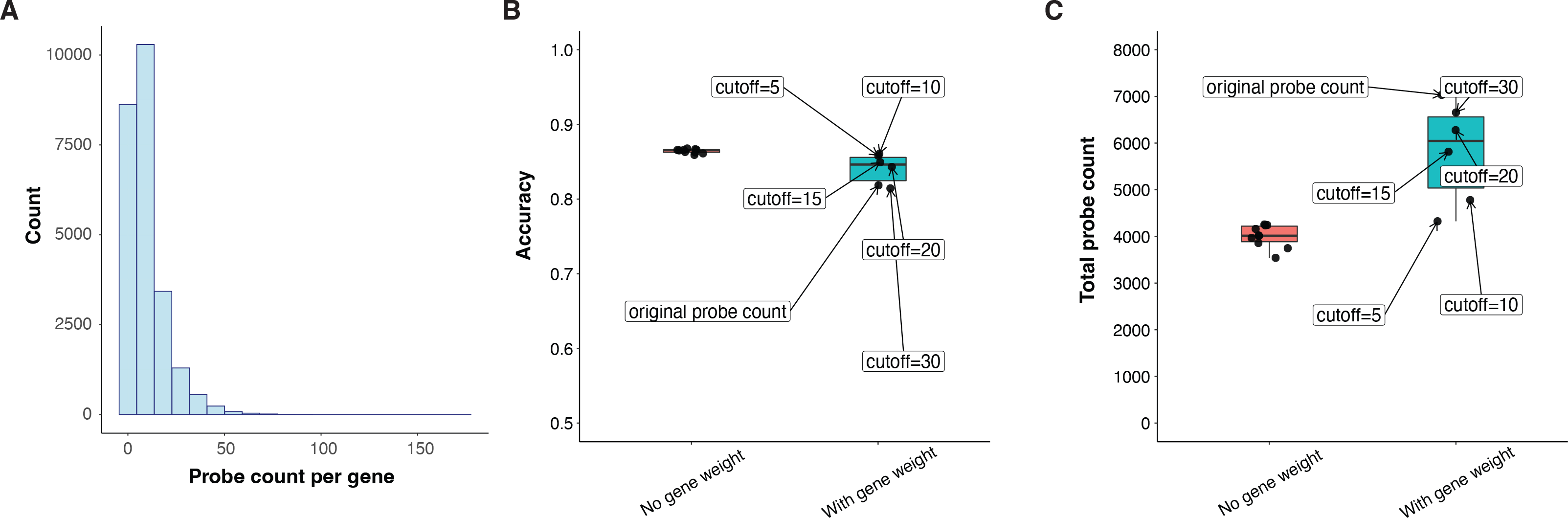
Weighted gene panel selection based on probe count per gene. **A**: Distribution of probe count per gene for the Zhang dataset. **B-C**: Distribution of accuracy (**B**) and total number of probes (**C**) of optimized gene panels from optimization without and with gene weight. Optimization without gene weight is performed 10 times. Optimization with gene weight is performed 6 times, each time with a different probe count cutoff (no cutoff, 5, 10, 15, 20, 30).

**Figure S7:**
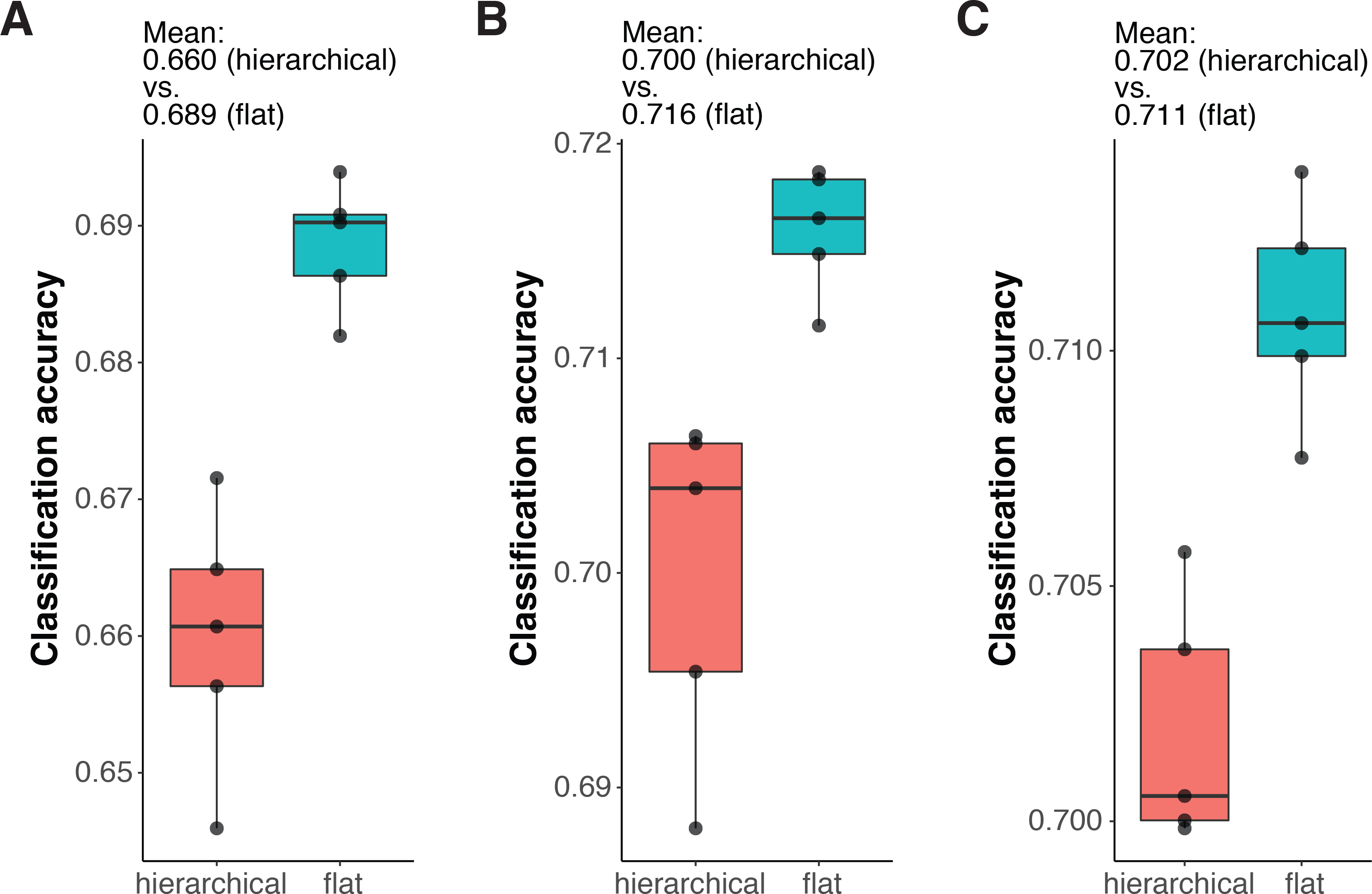
Accuracy of optimized gene panels using flat vs. hierarchical gene selection. **A-C**: Distribution of accuracy of optimized gene panels using flat vs. hierarchical gene selection for Moffit (**A**), Codeluppi (**B**), and Zhang (**C**), respectively. Both flat and hierarchical gene selection are performed 5 times.

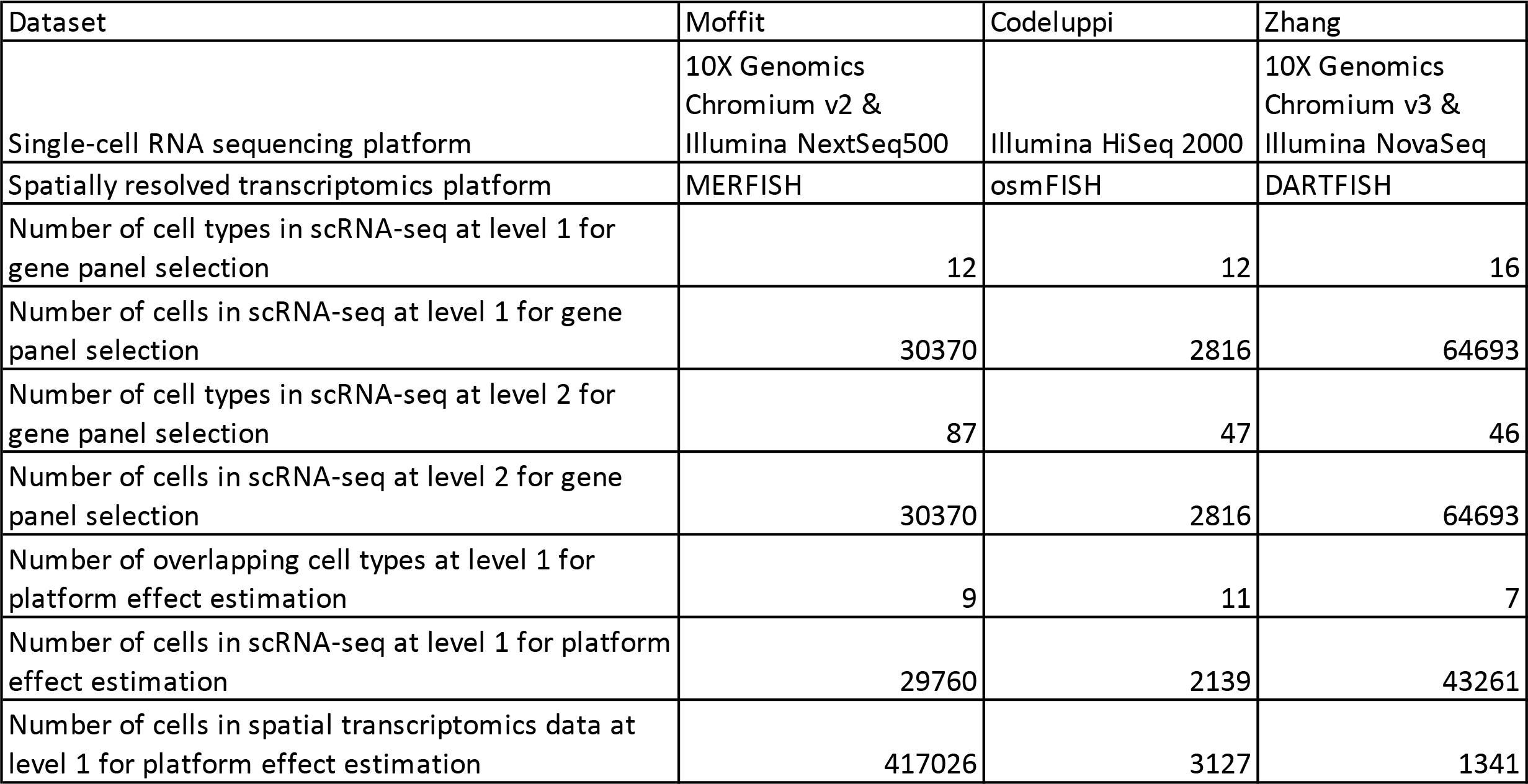
Information of the Moffit, Codeluppi, and Zhang dataset.

Information of the Moffit, Codeluppi, and Zhang dataset
| Dataset | Moffit | Codeluppi | Zhang |
| --- | --- | --- | --- |
| Single-cell RNA sequencing platform | 10X Genomics Chromium v2 & Illumina NextSeq500 | Illumina HiSeq 2000 | 10X Genomics Chromium v3 & Illumina NovaSeq |
| Spatially resolved transcriptomics platform | MERFISH | osmFISH | DARTFISH |
| Number of cell types in scRNA-seq at level 1 for gene panel selection | 12 | 12 | 16 |
| Number of cells in scRNA-seq at level 1 for gene panel selection | 30370 | 2816 | 64693 |
| Number of cell types in scRNA-seq at level 2 for gene panel selection | 87 | 47 | 46 |
| Number of cells in scRNA-seq at level 2 for gene panel selection | 30370 | 2816 | 64693 |
| Number of overlapping cell types at level 1 for platform effect estimation | 9 | 11 | 7 |
| Number of cells in scRNA-seq at level 1 for platform effect estimation | 29760 | 2139 | 43261 |
| Number of cells in spatial transcriptomics data at level 1 for platform effect estimation | 417026 | 3127 | 1341 |

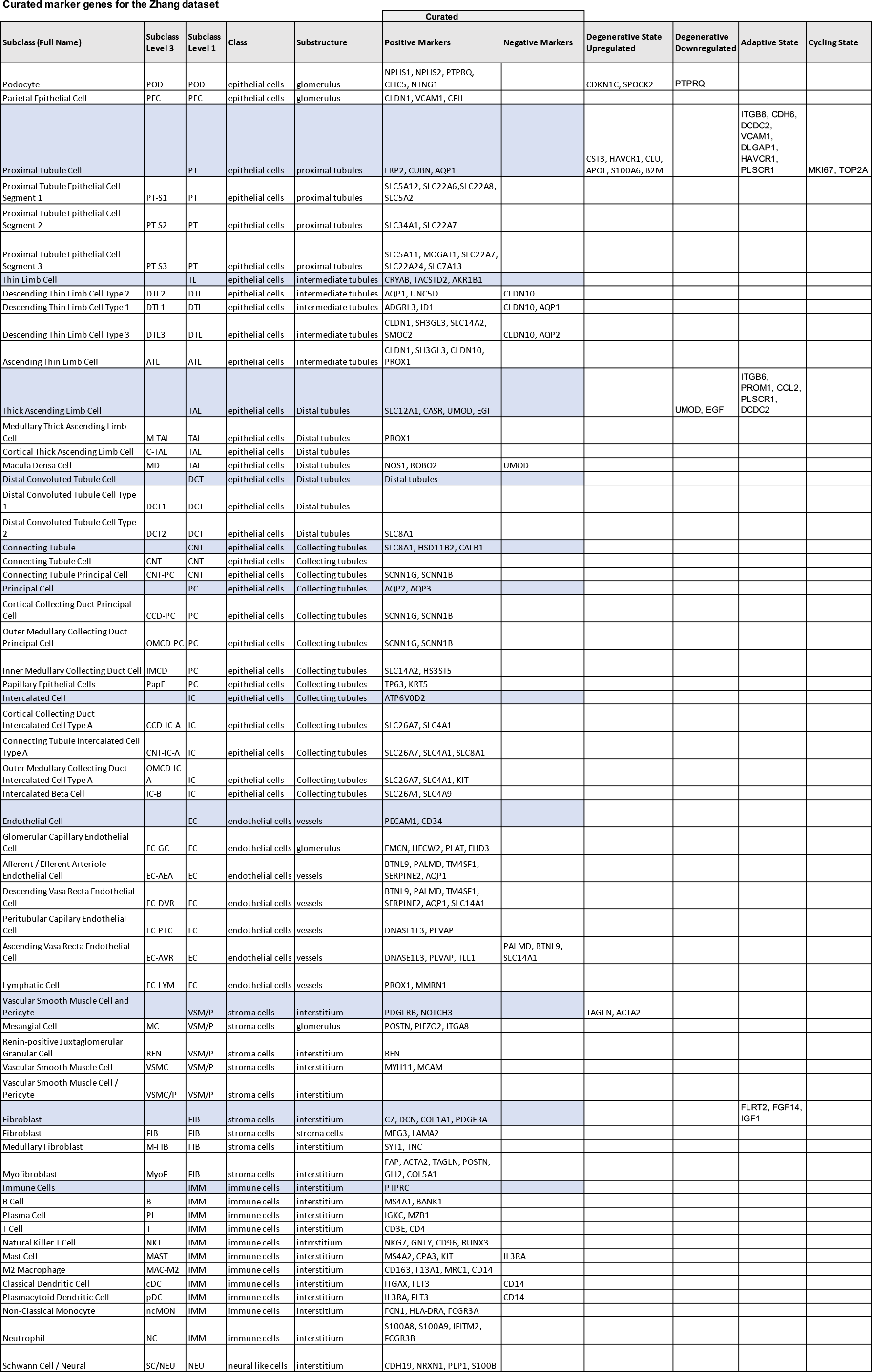
Curated marker genes for the Zhang dataset.

## Notes

https://doi.org/10.5281/zenodo.6946054

https://github.com/kharchenkolab/gpsFISH

### Summary of Updates

Updates were mainly made on the Discussion and Methods section for better clarity.

## Reference

1. Arendt D. The evolution of cell types in animals: emerging principles from molecular studies. Nat Rev Genet. 2008;9:868–82.

2. Elmentaite R, Domínguez Conde C, Yang L, Teichmann SA. Single-cell atlases: shared and tissue-specific cell types across human organs. Nat Rev Genet. 2022;23:395–410.

3. Lindeboom RGH, Regev A, Teichmann SA. Towards a Human Cell Atlas: Taking Notes from the Past. Trends in Genetics. 2021;37:625–30.

4. Zeng H. What is a cell type and how to define it? Cell. 2022;185:2739–55.

5. Kölsch Y, Hahn J, Sappington A, Stemmer M, Fernandes AM, Helmbrecht TO, et al. Molecular classification of zebrafish retinal ganglion cells links genes to cell types to behavior. Neuron. 2021;109:645–662.e9.

6. Osterhout JA, Kapoor V, Eichhorn SW, Vaughn E, Moore JD, Liu D, et al. A preoptic neuronal population controls fever and appetite during sickness. Nature. 2022;606:937–44.

7. Xu S, Yang H, Menon V, Lemire AL, Wang L, Henry FE, et al. Behavioral state coding by molecularly defined paraventricular hypothalamic cell type ensembles. Science. 2020;370:eabb2494.

8. Elmentaite R, Kumasaka N, Roberts K, Fleming A, Dann E, King HW, et al. Cells of the human intestinal tract mapped across space and time. Nature. 2021;597:250–5.

9. Armingol E, Officer A, Harismendy O, Lewis NE. Deciphering cell–cell interactions and communication from gene expression. Nat Rev Genet. 2021;22:71–88.

10. Chen W-T, Lu A, Craessaerts K, Pavie B, Sala Frigerio C, Corthout N, et al. Spatial Transcriptomics and In Situ Sequencing to Study Alzheimer’s Disease. Cell. 2020;182:976–991.e19.

11. Hwang WL, Jagadeesh KA, Guo JA, Hoffman HI, Yadollahpour P, Reeves JW, et al. Single-nucleus and spatial transcriptome profiling of pancreatic cancer identifies multicellular dynamics associated with neoadjuvant treatment. Nature Genetics [Internet]. 2022; Available from: https://doi.org/10.1038/s41588-022-01134-8

12. Darmanis S, Sloan SA, Zhang Y, Enge M, Caneda C, Shuer LM, et al. A survey of human brain transcriptome diversity at the single cell level. Proc Natl Acad Sci USA. 2015;112:7285–90.

13. Macosko EZ, Basu A, Satija R, Nemesh J, Shekhar K, Goldman M, et al. Highly Parallel Genome-wide Expression Profiling of Individual Cells Using Nanoliter Droplets. Cell. 2015;161:1202–14.

14. Tasic B, Menon V, Nguyen TN, Kim TK, Jarsky T, Yao Z, et al. Adult mouse cortical cell taxonomy revealed by single cell transcriptomics. Nat Neurosci. 2016;19:335– 46.

15. Ner-Gaon H, Melchior A, Golan N, Ben-Haim Y, Shay T. JingleBells: A Repository of Immune-Related Single-Cell RNA–Sequencing Datasets. JI. 2017;198:3375–9.

16. Trapnell C. Defining cell types and states with single-cell genomics. Genome Res. 2015;25:1491–8.

17. Regev A, Teichmann SA, Lander ES, Amit I, Benoist C, Birney E, et al. The Human Cell Atlas. eLife. 2017;6:e27041.

18. Marx V. Method of the Year: spatially resolved transcriptomics. Nat Methods. 2021;18:9–14.

19. Chen R, Blosser TR, Djekidel MN, Hao J, Bhattacherjee A, Chen W, et al. Decoding molecular and cellular heterogeneity of mouse nucleus accumbens. Nat Neurosci. 2021;24:1757–71.

20. Fernandez J. Molecular atlas of the adult mouse brain. SCIENCE ADVANCES. 2020;14.

21. Rao A, Barkley D, França GS, Yanai I. Exploring tissue architecture using spatial transcriptomics. Nature. 2021;596:211–20.

22. Zhang M, Eichhorn SW, Zingg B, Yao Z, Cotter K, Zeng H, et al. Spatially resolved cell atlas of the mouse primary motor cortex by MERFISH. Nature. 2021;598:137– 43.

23. Wang Y, Eddison M, Fleishman G, Weigert M, Xu S, Wang T, et al. EASI-FISH for thick tissue defines lateral hypothalamus spatio-molecular organization. Cell. 2021;184:6361–6377.e24.

24. Moffitt JR, Bambah-Mukku D, Eichhorn SW, Vaughn E, Shekhar K, Perez JD, et al. Molecular, spatial, and functional single-cell profiling of the hypothalamic preoptic region. Science. 2018;362:eaau5324.

25. Chen KH, Boettiger AN, Moffitt JR, Wang S, Zhuang X. Spatially resolved, highly multiplexed RNA profiling in single cells. Science. 2015;348:aaa6090–aaa6090.

26. Codeluppi S, Borm LE, Zeisel A, La Manno G, van Lunteren JA, Svensson CI, et al. Spatial organization of the somatosensory cortex revealed by osmFISH. Nat Methods. 2018;15:932–5.

27. Shah S, Lubeck E, Zhou W, Cai L. In Situ Transcription Profiling of Single Cells Reveals Spatial Organization of Cells in the Mouse Hippocampus. Neuron. 2016;92:342–57.

28. Cai M, Zhang K. Spatial mapping of single cells in human cerebral cortex using DARTFISH: A highly multiplexed method for in situ quantification of targeted RNA transcripts. eScholarship, University of California; 2019.

29. Wang X, Allen WE, Wright MA, Sylwestrak EL, Samusik N, Vesuna S, et al. Three-dimensional intact-tissue sequencing of single-cell transcriptional states. Science. 2018;361:eaat5691.

30. Chen X, Sun Y-C, Church GM, Lee JH, Zador AM. Efficient in situ barcode sequencing using padlock probe-based BaristaSeq. Nucleic Acids Research. 2018;46:e22–e22.

31. Chen X, Sun Y-C, Zhan H, Kebschull JM, Fischer S, Matho K, et al. High-Throughput Mapping of Long-Range Neuronal Projection Using In Situ Sequencing. Cell. 2019;179:772–786.e19.

32. Gyllborg D, Langseth CM, Qian X, Choi E, Salas SM, Hilscher MM, et al. Hybridization-based *in situ* sequencing (HybISS) for spatially resolved transcriptomics in human and mouse brain tissue. Nucleic Acids Research. 2020;48:e112–e112.

33. Chen A, Liao S, Cheng M, Ma K, Wu L, Lai Y, et al. Spatiotemporal transcriptomic atlas of mouse organogenesis using DNA nanoball-patterned arrays. Cell. 2022;185:1777–1792.e21.

34. Ståhl PL, Salmén F, Vickovic S, Lundmark A, Navarro JF, Magnusson J, et al. Visualization and analysis of gene expression in tissue sections by spatial transcriptomics. Science. 2016;353:78–82.

35. Rodriques SG, Stickels RR, Goeva A, Martin CA, Murray E, Vanderburg CR, et al. Slide-seq: A scalable technology for measuring genome-wide expression at high spatial resolution. 2019;6.

36. Vickovic S, Eraslan G, Salmén F, Klughammer J, Stenbeck L, Schapiro D, et al. High-definition spatial transcriptomics for in situ tissue profiling. Nat Methods. 2019;16:987–90.

37. Liu Y, Yang M, Deng Y, Su G, Enninful A, Guo CC, et al. High-Spatial-Resolution Multi-Omics Sequencing via Deterministic Barcoding in Tissue. Cell. 2020;183:1665–1681.e18.

38. Cho C-S, Xi J, Si Y, Park S-R, Hsu J-E, Kim M, et al. Microscopic examination of spatial transcriptome using Seq-Scope. Cell. 2021;184:3559–3572.e22.

39. Fu X, Sun L, Chen JY, Dong R, Lin Y, Palmiter RD, et al. Continuous Polony Gels for Tissue Mapping with High Resolution and RNA Capture Efficiency. bioRxiv. 2021;2021.03.17.435795.

40. Subramanian A, Narayan R, Corsello SM, Peck DD, Natoli TE, Lu X, et al. A Next Generation Connectivity Map: L1000 Platform and the First 1,000,000 Profiles. Cell. 2017;171:1437–1452.e17.

41. Missarova A, Jain J, Butler A, Ghazanfar S, Stuart T, Brusko M, et al. geneBasis: an iterative approach for unsupervised selection of targeted gene panels from scRNA-seq. Genome Biology. 2021;22:333.

42. Liang S, Mohanty V, Dou J, Miao Q, Huang Y, Müftüoğlu M, et al. Single-cell manifold-preserving feature selection for detecting rare cell populations. Nat Comput Sci. 2021;1:374–84.

43. Dumitrascu B, Villar S, Mixon DG, Engelhardt BE. Optimal marker gene selection for cell type discrimination in single cell analyses. Nat Commun. 2021;12:1186.

44. Vargo AHS, Gilbert AC. A rank-based marker selection method for high throughput scRNA-seq data. BMC Bioinformatics. 2020;21:477.

45. Aevermann BD, Zhang Y, Novotny M, Keshk M, Bakken TE, Miller JA, et al. A machine learning method for the discovery of minimum marker gene combinations for cell-type identification from single-cell RNA sequencing. Genome Res. 2021;gr.275569.121.

46. 46. Bakken TE, Hodge RD, Miller JA, Yao Z, Nguyen TN, Aevermann B, et al. Single-nucleus and single-cell transcriptomes compared in matched cortical cell types. Soriano E, editor. PLoS ONE. 2018;13:e0209648.

47. Cable DM, Murray E, Zou LS, Goeva A, Macosko EZ, Chen F, et al. Robust decomposition of cell type mixtures in spatial transcriptomics. Nat Biotechnol. 2022;40:517–26.

48. Okochi Y, Sakaguchi S, Nakae K, Kondo T, Naoki H. Model-based prediction of spatial gene expression via generative linear mapping. Nat Commun. 2021;12:3731.

49. Andersson A, Bergenstråhle J, Asp M, Bergenstråhle L, Jurek A, Fernández Navarro J, et al. Single-cell and spatial transcriptomics enables probabilistic inference of cell type topography. Commun Biol. 2020;3:565.

50. the FANTOM Consortium, Liang C, Forrest ARR, Wagner GP. The statistical geometry of transcriptome divergence in cell-type evolution and cancer. Nat Commun. 2015;6:6066.

51. Pliner HA, Shendure J, Trapnell C. Supervised classification enables rapid annotation of cell atlases. Nat Methods. 2019;16:983–6.

52. Tasic B. Single cell transcriptomics in neuroscience: cell classification and beyond. Current Opinion in Neurobiology. 2018;50:242–9.

53. Zeng H, Sanes JR. Neuronal cell-type classification: challenges, opportunities and the path forward. Nat Rev Neurosci. 2017;18:530–46.

54. Yuste R, Hawrylycz M, Aalling N, Aguilar-Valles A, Arendt D, Armañanzas R, et al. A community-based transcriptomics classification and nomenclature of neocortical cell types. Nat Neurosci. 2020;23:1456–68.

55. Bard J, Rhee SY, Ashburner M. An ontology for cell types. Genome Biology. 2005;5.

56. Bakken T, Cowell L, Aevermann BD, Novotny M, Hodge R, Miller JA, et al. Cell type discovery and representation in the era of high-content single cell phenotyping. BMC Bioinformatics. 2017;18:559.

57. Ashburner M, Ball CA, Blake JA, Botstein D, Butler H, Cherry JM, et al. Gene Ontology: tool for the unification of biology. Nat Genet. 2000;25:25–9.

58. The Gene Ontology Consortium, Carbon S, Douglass E, Good BM, Unni DR, Harris NL, et al. The Gene Ontology resource: enriching a GOld mine. Nucleic Acids Research. 2021;49:D325–34.

59. Zeisel A, Muñoz-Manchado AB, Codeluppi S, Lönnerberg P, La Manno G, Juréus A, et al. Cell types in the mouse cortex and hippocampus revealed by single-cell RNA-seq. Science. American Association for the Advancement of Science; 2015;347:1138–42.

60. Lake BB, Menon R, Winfree S, Hu Q, Ferreira RM, Kalhor K, et al. An atlas of healthy and injured cell states and niches in the human kidney. bioRxiv. 2021;2021.07.28.454201.

61. Goldberg DE. Genetic Algorithms in Search, Optimization and Machine Learning. 1st ed. USA: Addison-Wesley Longman Publishing Co., Inc.; 1989.

62. Holland JH. Adaptation in Natural and Artificial Systems: An Introductory Analysis with Applications to Biology, Control, and Artificial Intelligence [Internet]. The MIT Press; 1992 [cited 2022 Jul 30]. Available from: https://doi.org/10.7551/mitpress/1090.001.0001

63. Gene Expression Omnibus. www.ncbi.nlm.nih.gov/geo

64. Dryad. https://datadryad.org/stash/dataset/doi:10.5061/dryad.8t8s248

65. Zeisel A, Muñoz-Manchado AB, Codeluppi S, Lönnerberg P, La Manno G, Juréus A, et al. Single-cell analysis of mouse cortex. http://linnarssonlab.org/cortex/

66. Codeluppi S, Borm LE, Zeisel A, La Manno G, van Lunteren JA, Svensson CI, et al. osmFISH: Spatial organization of the somatosensory cortex revealed by cyclic smFISH. http://linnarssonlab.org/osmFISH/

67. Zhang Y, Petukhov V, Biederstedt E, Que R, Zhang K, Kharchenko PV. gpsFISH analysis code and data (Zenodo link). 2023; https://doi.org/10.5281/zenodo.7613712

68. ppDesigner: Algorithm to design Padlock Probes. http://genome-tech.ucsd.edu/public/Gen2_BSPP/ppDesigner/ppDesigner.php

69. Zhang Y, Petukhov V, Biederstedt E, Que R, Zhang K, Kharchenko PV. gpsFISH R package. GitHub. 2023; https://github.com/kharchenkolab/gpsFISH

70. Zhang Y, Petukhov V, Biederstedt E, Que R, Zhang K, Kharchenko PV. Pre-fitted Bayesian models (Zenodo link). 2023; https://doi.org/10.5281/zenodo.6946054

